# NKp30 and NKG2D contribute to natural killer recognition of HIV-infected cells

**DOI:** 10.1101/2024.06.24.600449

**Authors:** Nancy Q. Zhao, Ruoxi Pi, David N. Nguyen, Thanmayi Ranganath, Christof Seiler, Susan Holmes, Alexander Marson, Catherine A. Blish

## Abstract

Natural killer (NK) cells respond rapidly in early HIV-1 infection. HIV-1 prevention and control strategies harnessing NK cells could be enabled by mechanistic understanding of how NK cells recognize HIV-infected T cells. Here, we profiled the phenotype of human primary NK cells responsive to autologous HIV-1-infected CD4^+^ T cells in vitro. We characterized the patterns of NK cell ligand expression on CD4^+^ T cells at baseline and after infection with a panel of transmitted/founder HIV-1 strains to identify key receptor-ligand pairings. CRISPR editing of CD4^+^ T cells to knockout the NKp30 ligand B7-H6, or the NKG2D ligands MICB or ULBP2 reduced NK cell responses to HIV-infected cells in some donors. In contrast, overexpression of NKp30 or NKG2D in NK cells enhanced their targeting of HIV-infected cells. Collectively, we identified receptor-ligand pairs including NKp30:B7-H6 and NKG2D:MICB/ULBP2 that contribute to NK cell recognition of HIV-infected cells.

## INTRODUCTION

Human immunodeficiency virus (HIV) remains a significant global public health issue, with 39 million people still living with HIV in 2022 and an estimated 1.3 million new infections yearly highlighting the need for improved strategies to prevent infection.^1^ Natural killer (NK) cells are innate immune cells that are among the earliest responders to viral infection. They rapidly expand during the earliest stages of HIV infection^2,3^, and have been implicated in both acquisition and immune control of HIV. Highly exposed seronegative individuals have increased constitutive NK cell activity^4,5^, suggesting that NK cell activity can protect against infection. Further, the number of CXCR5^+^ NK cells in B cell follicles of secondary lymphoid tissue (SLT) inversely correlated with the viral load of HIV, simian immunodeficiency virus (SIV), or SHIV/HIV hybrid virus (SHIV) during chronic infections.^6–8^ Higher functionality of CXCR5^+^ NK cells indicated by CD107a or IFNγ expression compared to CXCR5^-^ counterparts suggests that NK cells may limit virus spread in T follicular helper cells (T_FH_), one of the cell types that naturally harbor persistent reservoirs of latent HIV.^6–8^ Finally, depletion of NK cells led to elevated HIV viral load in plasma, lymph nodes, the spleen and liver in humanized mice, indicating the role of NK cells in suppressing HIV infection.^9^ Improved understanding of the mechanisms NK cells utilize to recognize and respond to HIV-1 infection could allow these immune functions to be harnessed to prevent, control, or even cure infection.

Much effort on NK cell recognition mechanisms have focused on interactions between HLA class I molecules on HIV-infected cells and the killer immunoglobulin receptors (KIRs) on NK cells. This is particularly relevant as multiple studies have demonstrated the importance of the KIR3DL1–HLA-B allotype interaction in slower disease progression to AIDS and lower viral load^10^, reduced risk of HIV infection in HIV-exposed seronegative subjects^11^, and improved control of HIV replication *in vitro*.^12,13^ HIV-1 infection induces downregulation of expression of the classical HLA class I molecules HLA-A, HLA-B, and HLA-C on infected cells^14–16^, which may be of help in the evasion of cytotoxic T lymphocyte responses but can instead trigger recognition by educated NK cells. Consistent with the idea that educated NK cells can play a critical role in HIV-1 control, high expression of both surface KIR3DL1 and its HLA ligand (the Bw4 epitope) leads to the generation of more potently reactive NK cells to HIV-infected cells^17^, presumably via stronger education and hence reactivity to ‘missing self’. In addition, NK cell education via KIRs determines their ability to respond to HIV-mediated downregulation of their cognate HLA class I ligand on target cells *in vitro*.^18^

Besides the interaction between HLA class I molecules and KIRs, there is growing evidence that HLA-E, a nonclassical HLA class I molecule, which interacts with NK cell receptors CD94/NKG2A and CD94/NKG2C, can regulate NK cell function in response to HIV-infected cells. Multiple HIV-derived peptides, when presented by HLA-E, have been identified to induce elevated functionality of NK cells, including enhanced cytotoxicity, degranulation or TNF-α secretion.^19–21^ It has been hypothesized that peptides binding to HLA-E can either interrupt the inhibitory interaction between HLA-E and CD94/NKG2A, or facilitate NK cell activation mediated by CD94/NKG2C. Nef of multiple HIV strains downregulates HLA-E in primary CD4^+^ T cells^22^, potentially influencing peptide presentation by HLA-E and interaction between HLA-E and its receptors.

In addition to recognition of HLA class I molecules, infected cells can also modulate the expression of ligands for other NK cell receptors, mediated mostly by HIV-1 accessory proteins. For example, Vpr is known to upregulate NKG2D ligands, particularly ULBP2.^23^ Conversely, Vpu and Nef downregulate these ligands, providing an avenue for immune escape.^24^ Vpu also downregulates NTB-A, a homotypic co-activating signaling lymphocytic activation molecule (SLAM) family receptor found on both CD4^+^ T cells as well as NK cells, and this has been reported to prevent NK cell activation against infected cells.^24–26^ Vpu of multiple HIV strains downregulates CD48, the ligand of another SLAM family receptor, CD244.^26^ Impairment of NTB-A-NTB-A and CD48-CD244 interactions dampens antibody-dependent cellular cytotoxicity (ADCC) towards HIV-infected primary CD4^+^ T cells.^26^ However, as NK cells integrate signaling from all their surface receptors to determine activation, it remains unclear whether the overall ligand modulation on infected cells leads to NK cell susceptibility or escape, and which receptor-ligand interactions are most prominently involved. The recent use of high-dimensional mass cytometry has allowed the simultaneous profiling of the numerous NK cell surface receptors and their respective ligands on infected cells^27,28^, providing a powerful tool to address overall pathogen-driven patterns of ligand modulation.

While the traditional paradigm suggests that innate immune cells are activated by invariant pathogen signatures and do not possess specificity, recent work has shown that this is not always the case. Indeed, a pandemic influenza H1N1 strain differentially modulates NK cell ligands and triggers different levels of IFN-D production by NK cells than a co-circulating H3N2 strain.^29^ As such, variability between viral strains that leads to different pathways of NK ligand modulation may lead to strain-specific responses by the innate immune system. HIV is extremely genetically diverse -even more so than influenza^30^ - and hence it remains unclear if particular strains may have different patterns of ligand modulation or induce differential recognition by NK cells.

Here, we use mass cytometry to simultaneously profile the expression of known NK cell ligands on HIV-infected CD4^+^ T cells, as well as the phenotypes of NK cells responding to autologous HIV-infected cells. We analyzed NK ligand expression on CD4^+^ T cells infected with a panel of HIV strains comprising transmitted/founder (T/F) and laboratory-adapted subtype B strains and an early subtype A strain to determine if patterns of ligand expression were differentially modulated by different strains. We used these analyses to determine a set of candidate receptor-ligand interactions that mediate NK cell recognition of HIV infection. Knockout of these T cell surface ligands individually with CRISPR editing, and comparison between NK cell response to HIV-infected and to mock-infected CD4^+^ T cells, suggested a potential role for B7-H6 (a ligand for NKp30), major histocompatibility complex (MHC) class I chain-related protein B (MICB, a NKG2D ligand), and UL16 binding protein 2 (ULBP2, another NKG2D ligand) in HIV-specific recognition by NK cells. We overexpressed NKp30 and NKG2D with mRNA transfection and confirmed their role of mediating NK cell recognition of HIV-infected and mock-infected CD4^+^ T cells.

## RESULTS

### An *in vitro* autologous co-culture system allows the detection of NK cell responses against HIV

We first optimized an *in vitro* co-culture system to allow the detection of HIV-specific responses, using CD4^+^ T cells infected *in vitro* with the subtype A HIV-1 virus Q23-17 (shortened as Q23) co-cultured with autologous NK cells.^31^ We observed consistent NK cell responses with both cytolytic degranulation (by the expression of the degranulation marker CD107a; Figure 1A) as well as cytokine production (for the canonical NK cell cytokine IFN-D; Figure 1B). Notably, healthy donors differed considerably in their functional response to autologous, HIV-infected CD4^+^ T cells. Both degranulation and cytokine production scaled negatively with increasing effector:target (E:T) ratio. In addition, we observed effective viral suppression mediated by NK cells with a reduction in levels of p24^+^ cells in the presence of NK cells; viral suppression scaled positively with E:T ratio, as expected (Figure 1C). As such, we were able to further utilize this *in vitro* system to probe specific interactions involved in NK cell recognition of HIV and the initiation of downstream HIV-specific responses.

**Figure 1:**
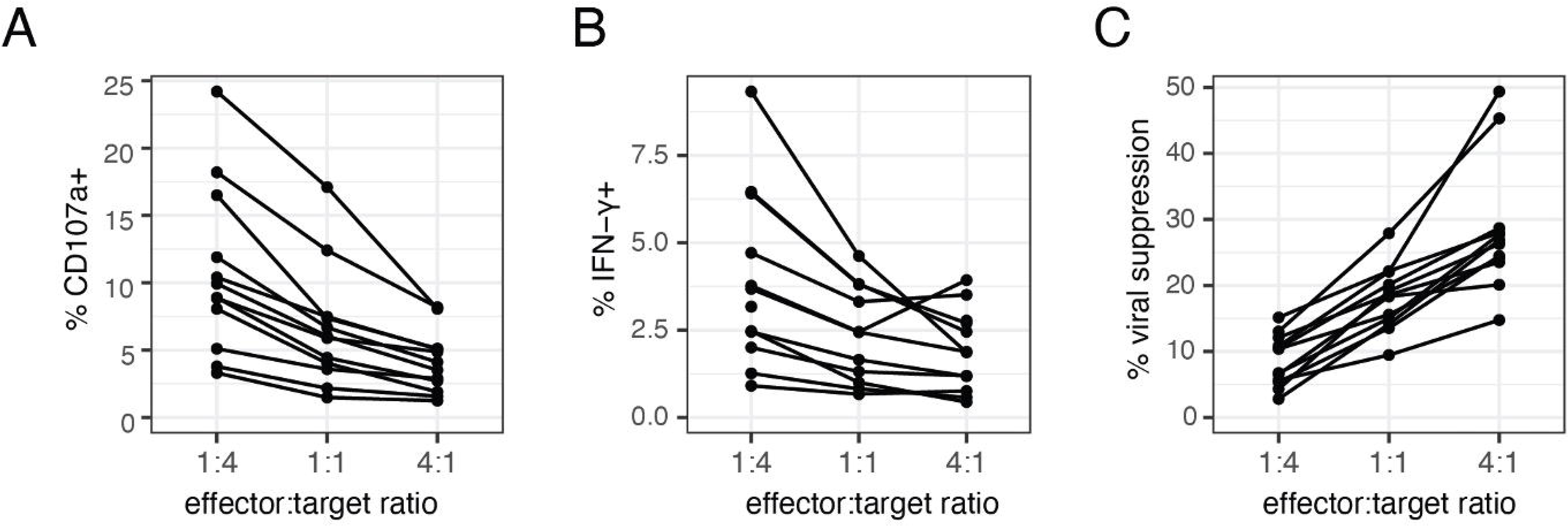
*An in vitro NK-CD4^+^ T cell co-culture system allows the dissection of responses to HIV-infected cells.* An *in vitro* co-culture system was used to detect NK cell responses to autologous CD4^+^ T cells infected with HIV strain Q23. Summary data for NK cell responses, for **(A)** cytolytic degranulation by the marker CD107a, **(B)** IFN-D production, and **(C)** viral suppression, at varying E:T ratios for 12 healthy donors tested in this assay. Viral suppression was calculated as the percent decrease in CD4^+^ T cells positive for the HIV Gag antigen p24 in the presence vs absence of NK cells.

### Phenotypic profiling of HIV-responsive NK cells with mass cytometry

To determine the receptors involved in triggering the NK cell functional responses observed, we used mass cytometry to simultaneously profile expression of NK cell surface receptors and functional markers in our co-culture system. We reasoned that cells performing any of the canonical NK cell functions, including cytotoxic activity (indirectly measured by CD107a induction) and cytokine secretion (IFN-D and TNF-D) could contribute to the suppression of HIV-1 replication in infected CD4^+^ T cells. Thus, for each sample we gated on NK cells that were positive for CD107a, IFN-D, and TNF-D (Figures S1A, B), and used Boolean gating to identify functionally responding cells (termed functional^+^, positive for *any* of the functional markers above), or non-functionally responding cells (termed functional^-^, negative for *all* of the markers above). To identify NK cell receptors that may contribute to the response, we then used a generalized linear mixed model (GLMM) implemented through CytoGLMM to compare paired functional^+^ and functional^-^ NK cells in co-culture with HIV-infected cells from each donor (Figure 2A).^32^ This revealed a distinct phenotype of responding functional cells; with the exception of 5 receptors (NKG2D, CD2, NTB-A, KIR2DS4 and PD-1), every NK receptor profiled was a significant predictor of either a functional^+^ or functional^-^ phenotype (Figure 2A). The top three predictors of responding functional^+^ cells were TIGIT, NKp30 and KIR3DL1, whereas the top three predictors of non-responding functional^-^ cells were CD16, NKp46, and CD62L. Importantly, when the same analysis was performed with NK cells co-cultured with mock-infected CD4^+^ T cells, we found that many of the predictors of responding cells were similar (Figure 2B), suggesting that the responding NK cell phenotype identified was not unique to infection, likely because the activation of CD4^+^ T cells required to infect them can upregulate stress ligands and induce NK cell killing.^33^ Indeed, when we compared functional cells from NK cells co-cultured with mock-infected cells, to functional^+^ cells with NK cells co-cultured with HIV-infected cells, we found few receptors that could predict functional status (Figure 2C). These results suggest that HIV-targeting by NK cells may not necessarily utilize distinct receptors but may instead rely on altered levels of signaling from these receptors.

**Figure 2:**
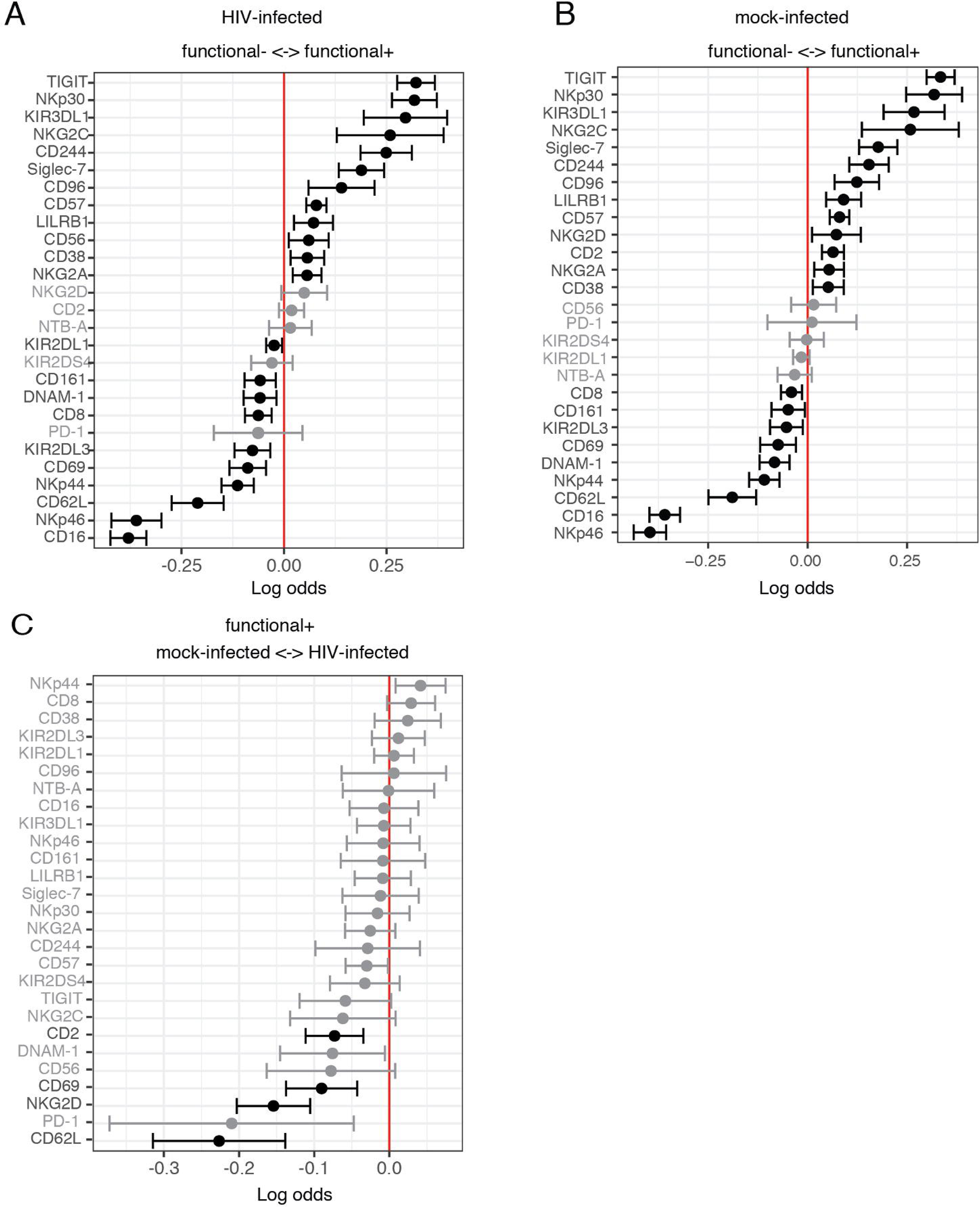
*Mass cytometry profiling of NK cell receptors and functional markers identifies the phenotypic features of responding NK cells* **(A, B)** A generalized linear mixed model (GLMM) with paired comparison was used to compare non-responding (functional^-^) and responding NK cells (functional^+^), after NK cells were co-cultured with HIV-infected (A) or mock-infected (B) CD4^+^ T cells from the same individual (n=20) at effector:target ratio of 1:4. **(C)** A GLMM with paired comparison was used to compare responding cells from NK cells in co-culture with either mock-infected or HIV-infected CD4^+^ T cells (n=20). Markers with adjusted p-value < 0.05 are shown in black and markers with adjusted p-value > 0.05 are shown in grey; adjusted *p*-values calculated on GLMM using Benjamini-Hochberg method to control the false discovery rate.

### HIV infection of CD4^+^ T cells induces upregulation of activating NK cell ligands

The presence of certain receptors on NK cells responding to HIV infection may merely represent markers of generally more functional NK cells, instead of receptors involved in the specific response to HIV infection. Indeed, the top predictor of responding cells was TIGIT (Figure 2A), which we have reported to mark a more functional NK cell subset.^34^ However, it is still under discussion if TIGIT influences NK cell response that is specific to HIV infection in primary CD4^+^ T cells.^34–37^ To gain insight into which NK cell ligands on primary CD4^+^ T cells are modulated by HIV infection and thereby trigger HIV-specific NK cell responses, we complemented NK receptor profiling with an analysis of NK cell ligand expression on HIV-infected CD4^+^ T cells. We utilized a CyTOF panel that included a set of known activating and inhibitory ligands for NK cell receptors to profile CD4^+^ T cells (gating strategy in Figure S1C) 24 hours after the inoculation of Q23 strain (HIV-infected in Figure 3A), as well as CD4^+^ T cells that were not inoculated with HIV but treated in parallel otherwise (mock-infected in Figure 3A). We also identified actively infected (HIV Gag p24^+^) and bystander (p24^-^) cells in the CD4^+^ T cell culture with HIV (Figures 3A). To understand the alterations in NK ligand expression in HIV-infected cells, we used *CytoGLMM* to identify surface ligands that predicted CD4^+^ T cells in the culture with HIV compared to mock-infected cells (Figure 3B).

**Figure 3:**
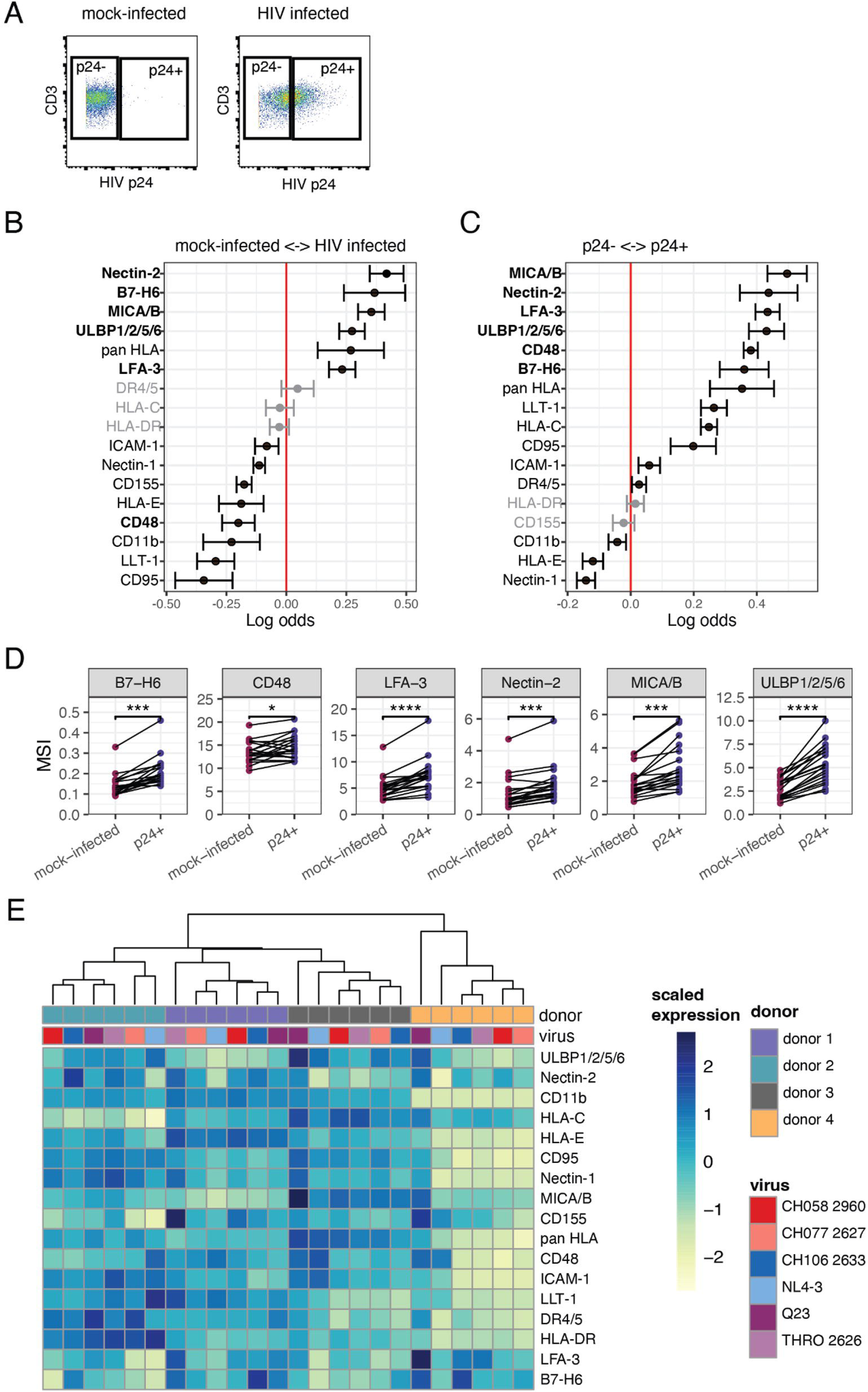
*A group of activating NK cell ligands are upregulated in HIV-infected cells.* Expression of NK cell ligands on CD4^+^ T cells was profiled by CyTOF. **(A)** Representative dot plots from mass cytometry, demonstrating the gating strategy to identify HIV-infected cells as determined by positive staining for HIV p24. A generalized linear mixed model (GLMM) with paired comparison was used to compare **(B)** mock-infected and HIV-infected cells (n=20), and **(C)** p24^-^ and p24^+^ cells from within infected wells (n=20). Markers with adjusted p-value < 0.05 are shown in black and markers with adjusted p-value > 0.05 are shown in grey; adjusted *p*-values calculated on GLMM using Benjamini-Hochberg method to control the false discovery rate. **(D)** Comparison of mean signal intensity (MSI) between mock-infected and p24^+^ cells for ligands of interest; individual donors are joined by a line. **(E)** Dendrogram shows results of unsupervised clustering based on mean asinh-transformed expression of each marker for each sample (4 donors, 6 HIV strains in total); heatmap shows scaled mean marker intensity in each sample, scaled within each marker to enable visualization of variation in expression of each NK ligand.

Furthermore, in the CD4^+^ T cell culture inoculated with HIV, we also identified predictors of p24^+^ cells compared to bystander p24^-^ T cells (Figure 3C). We found that Nectin-2 (ligand of DNAM-1), B7-H6 (ligand of NKp30), LFA-3 (ligand of CD2), MICA/B (ligands of NKG2D), and ULBP1/2/5/6 (ligands of NKG2D) were all predictors for infected cells (Figures 3B and C); CD48 (ligand of CD244) was a predictor of p24^+^ cells when compared to p24^-^ bystanders (Figure 3C), although not predictive of total CD4^+^ T cells in the culture with HIV compared to mock-infected controls (Figure 3B). The increased expression of these markers was confirmed by analyzing mean signal intensity (MSI), which showed that MSI was higher in p24^+^ cells compared to mock-infected cells for all these ligands (Figure 3D). Notably, these are all activating ligands for NK cells through interaction with their receptors; the expression of the corresponding receptors on NK cells, including NKp30 (receptor for B7-H6) and CD244 (receptor for CD48), was also higher on functionally responding NK cells (Figure 2A) revealing potential pathways for NK cell recognition of HIV-infected cells. Thus, using a combination of CyTOF screens on NK cell receptors and on their ligands expressed by CD4^+^ T cells, we identified a set of 6 candidate receptor-ligand pairs that had a potential role in NK cell recognition of HIV-infected cells: DNAM-1:Nectin-2, NKp30:B7-H6, CD2:LFA-3, CD244:CD48, NKG2D:MICA/B, and NKG2D:ULBP1/2/5/6.

### The pattern of NK cell ligand expression in HIV-infected CD4^+^ T cells varies with different virus strains and among individuals

To further understand whether different strains of HIV might induce variable patterns of NK ligand alteration therefore differentially influence NK cell recognition, we compared ligand expression on CD4^+^ T cells infected with each of 6 HIV strains, using the same ligand panel for CyTOF analysis as above.

These strains included Q23, a commonly used lab-adapted strain NL4-3, as well as a panel of T/F strains, which are particularly relevant for the study of early innate immune cell responses. For each HIV strain, a multiplicity of infection (MOI) was chosen to achieve 40-80% of total cells positively staining for HIV p24 (Figure S2A). To look at overall NK ligand expression patterns on infected cells (p24^+^) only, we used a PCA plot to visualize all samples colored by virus strain (Figure S2B top left panel) and blood donor (Figure S2B bottom left panel). We found that samples grouped predominantly based on blood donor (Figure S2B) suggesting a greater impact of blood donor than of virus strain on the overall pattern of NK cell ligand expression. To better understand the differences in NK ligand modulation driven by donor variability, compared to driven by different viral strains, we performed unsupervised hierarchical clustering based on mean marker expression for all profiled ligands in each sample (Figure 3E). We again found that samples clustered by donor but not viral strain, reinforcing that patterns of NK ligand expression are driven predominantly by donor-to-donor variance. To verify whether ligands upregulated in Q23 infection were shared across other HIV strains, we compared individual ligand expression across all the 6 HIV strains. CD48, LFA-3 and MICA/B were differentially regulated by different strains while the upregulation of B7-H6, ULBP1/2/5/6 and Nectin-2 in the infected CD4^+^ T cells are conserved among Q23 and all 4 T/F strains in most of the donors (Figure S2C). Expression levels of MICA/B and ULBP1/2/5/6 were higher in CD4^+^ T cells infected with Q23 than with all the subtype B T/F strains in most of the donors (Figure S2C). Collectively, our data suggest that NK cell ligand expression can be differentially shaped by different HIV strains yet highly variable among individuals.

### Knockout of B7-H6 and ligands of NKG2D diminish the HIV-specific NK cell response

To assess the individual roles of each of these ligands, we knocked each of them out separately in CD4^+^ T cells, and then probed the NK cell response in the presence or absence of HIV infection (Figure 4A). CD4^+^ T cells were electroporated with CRISPR/Cas9 ribonucleoproteins (RNPs) containing a pool of three different guide RNAs (gRNAs) against each ligand of interest. We also included non-treated controls (NT), which were not electroporated, as well as a control targeting AAVS1, a ‘safe harbor’ region of the genome. Targeting of AAVS1 did not lead to any changes in ligand expression compared to NT controls, indicating that RNP electroporation or targeted double-strand breaks in the genome did not lead to any modulation of ligand expression (Figure S3A). To test for the efficiency of ligand knockout, we examined surface expression of each ligand 6 days post-electroporation by flow cytometry, in both mock-and HIV-infected cells. Despite considerable variability between donors in the level of NK cell ligand expression both before and after knockout, we noted an expected reduction in the expression level of most ligands in targeted knockouts compared to NT in total CD4^+^ T cells (Figures 4B and S3B) within each donor. The reduction in B7-H6, Nectin-2, MICA, MICB, ULBP1, and ULBP2 was mild in some or all the donors compared to CD48 and LFA-3, possibly due to their low level of expression even in non-treated CD4^+^ T cells (Figures 4B and S3B). CRISPR editing did not lead to a discernible effect on T cell susceptibility to HIV infection (data not shown).

**Figure 4:**
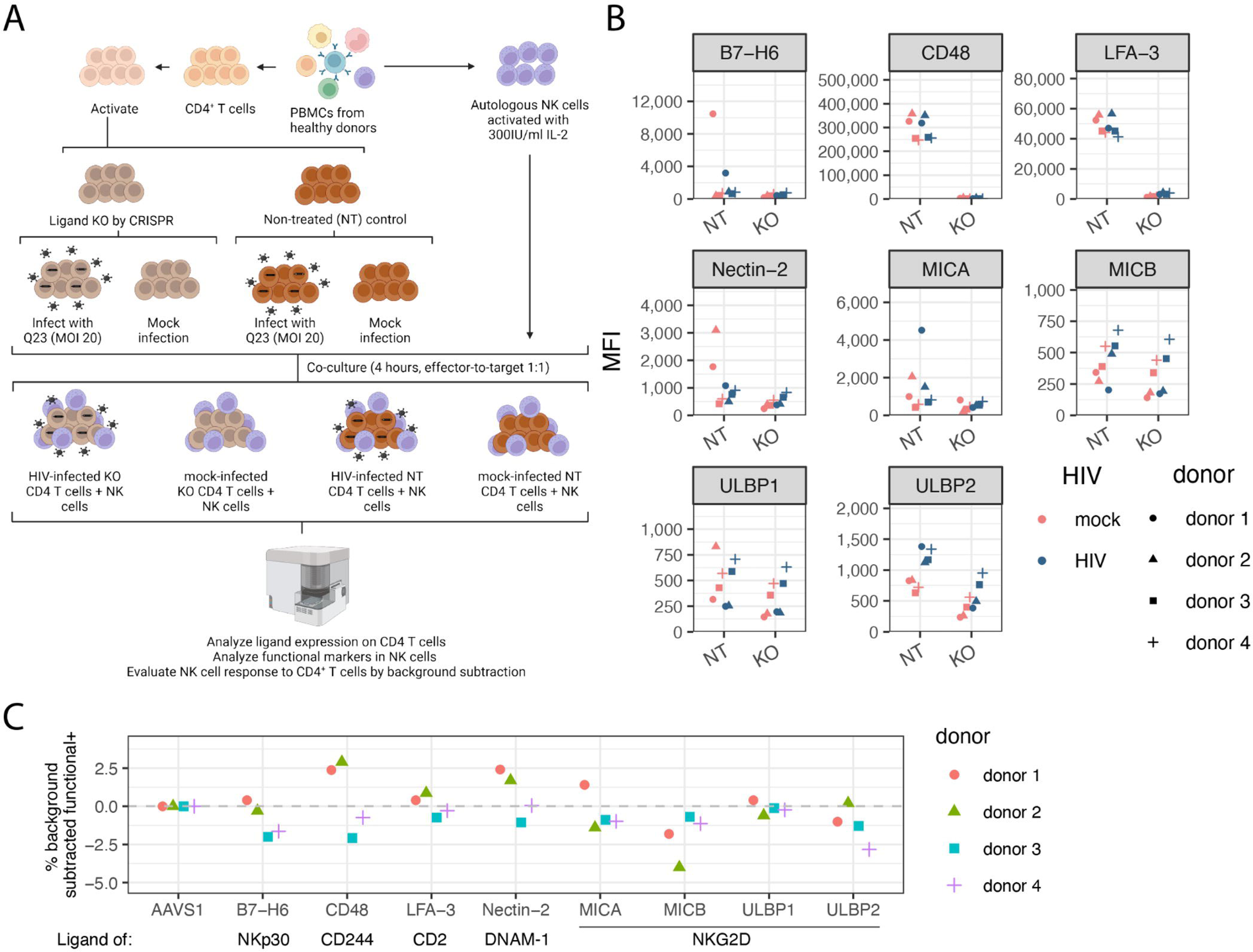
*Knockout of T cell targets reveals that reduction of B7-H6 or ligands of NKG2D reduces the HIV-specific NK cell response.* **(A)** A schematic of the experimental design for testing the function of NK cells in response to CD4^+^ T cells in which individual NK cell ligands were knocked out with CRISPR. This image was created with BioRender.com. **(B)** Mean fluorescence intensity (MFI) of each ligand tested, as well as non-treated (NT) control, in mock-(in pink) and HIV-infected (in blue) CD4^+^ T cells. Each donor is indicated by a different symbol. **(C)** Quantification of the HIV-specific effect of knocking out each ligand within each donor. For each ligand knockout, we determined the HIV-specific response by subtracting the percentage of functional^+^ NK cells from the mock-infected CD4^+^ T cells from the percentage of functional^+^ NK cells from the HIV-infected CD4^+^ T cells. We then normalized to the HIV-specific response for each donor to the control AAVS1-edited group by subtracting that level, such that negative values indicate a diminished functional response and positive values indicate an enhanced functional response compared to the control AAVS1-edited group. n=4.

To identify the effects of these ligand knockouts on NK cell responses in the context of HIV infection, we co-cultured edited CD4^+^ T cells that were either mock-infected or Q23-infected with autologous NK cells, and then monitored the NK cell functional response using flow cytometry (Figure 4A). We used the frequency of functional NK cells as defined above (% functional^+^) to evaluate overall NK cell responses. Unexpectedly, in some donors, knockout of CD48, MICA, and MICB, led to increased functional responses compared to NT controls in response to both mock- and HIV-infected cells (Figure S3C). As we were predominantly interested in HIV-specific responses, we applied mock-subtraction to identify HIV-specific responses, and donor-level normalization to evaluate the effect of knocking out each NK cell ligand within each donor by comparing them to the control group where AAVS1 was edited (Figure 4C). Despite the low level of expression in non-treated CD4^+^ T cells, knockout of B7-H6 (NKp30 ligand) and ligands of NKG2D, including MICB, and ULBP2 led to a reduction in the HIV-specific response to variable extents in most of the donors tested. However, knockout of any of the ligands did not lead to a complete abrogation of HIV-specific NK cell targeting (Figure S3C). As such, while knockout of B7-H6 and ligands of NKG2D diminished the NK cell response towards HIV-infected CD4^+^ T cells from some donors, knockout of any individual NK cell ligand did not fully abolish the HIV-specific response.

### Overexpression of NKp30 and NKG2D enhances NK cell response to HIV-infected CD4^+^ T cells

Since all HIV strains tested led to upregulation of ULBP1/2/5/6 and B7-H6 on HIV-infected CD4^+^ T cells, and the knock-out of B7-H6, MICB and ULBP2 modestly reduced NK cell responses to infected cells in most donors, we further investigated if their receptors, NKp30 and NKG2D, influence NK cell response. We also investigated the role of CD244 (2B4) in NK cell response to CD4^+^ T cells since CD244 is among the top predictors of functional NK cells in the co-culture with both Q23-infected and mock-infected CD4^+^ T cells (Figures 2A, B) yet knockout of its ligand, CD48, led to variable effects on the HIV-specific NK cell response in different donors (Figure 4C). We overexpressed NKp30, NKG2D, and CD244 in NK cells with charge-altering releasable transporter (CART) transfection. CART transfection allows highly efficient delivery and expression of exogenous mRNA in human primary NK cells, preserving their viability and causing minimal impact on their phenotypes.^38,39^ We isolated NK cells from peripheral blood mononuclear cells (PBMCs) of healthy donors and transfected them with CART and the encoding mRNAs of NKp30, NKG2D and CD244. To distinguish the NK cells that overexpress these receptors after transfection and the ones that were not transfected but naturally express the receptors, we co-transfected the NK cells with GFP mRNA to indicate the translation of exogenous mRNAs. Since DAP10 is required for the surface expression of human NKG2D and serves as an signaling adaptor of NKG2D^40,41^, we first ascertained if excessive DAP10 mRNA was required for surface expression of exogenous NKG2D in human NK cells. Interestingly, the endogenous level of DAP10 in NK cells is not enough to support the surface expression of the exogenous NKG2D (Figure S4). Co-transfection of NKG2D and DAP10 mRNAs at a ratio of 3:1 maximizes the expression level of NKG2D in NKG2D^+^GFP^+^ double positive NK cells (Figure S4C).

We evaluated if NK cell function in response to Q23-infected CD4^+^ T cells is altered with the overexpression of NKp30, NKG2D, or CD244 (Figure 5A). At 16-19 hours after NK cells were transfected with each of the three combinations of encoding mRNA, including NKp30+GFP, NKG2D+DAP10+GFP, and CD244+GFP, we observed a population of GFP^+^ NK cells (30-60%, Figures 5B and C) that expressed higher levels of NKp30, NKG2D, or CD244, respectively, compared to the GFP^-^ NK cells in the same well (Figure 5B). We co-cultured transfected NK cells (including both GFP^-^ and GFP^+^ in the same well) in each of the three groups mentioned above with autologous Q23-infected or mock-infected CD4^+^ T cells at E:T ratio of 1:4 for 4 hours. NK cells that were cultured alone serve as controls that indicate the background level of NK cell function. When NK cells were co-transfected with combinations of mRNA that encode either NKp30+GFP or NKG2D+DAP10+GFP, the percentage of functional^+^ NK cells (as defined above, gating strategy in figure S5) in the GFP^+^ population was significantly higher than in the GFP^-^ population in the same well in the presence of Q23-infected CD4^+^ T cells (Figure 5D top panel). When NKs were co-transfected with CD244 and GFP, the percentage of functional^+^ in the GFP^+^ population was significantly reduced compared to the GFP^-^ population (Figure 5D top panel). The alteration of function in GFP^+^ NK cells is not due to the expression of GFP or excessive amount of DAP10 because transfection of GFP mRNA alone or co-transfection with DAP10 and GFP only marginally influenced the percentage of functional^+^ NK cells (Figure 5D top panel). Furthermore, when we stratified NK cells based on the expression level of NKp30, NKG2D, or CD244 to high, intermediate and low/negative (Figure S6A), we noticed positive correlations between the percentage of functional^+^ NK cells and the expression level of NKp30 or NKG2D (Figures S6B and C, top panel). We also observed that the percentage of functional^+^ NK cells in the CD244^high^ population was reduced compared to the CD244^int^ and CD244^low/-^ populations (Figure S6D, top panel). However, the alteration of NK cell function due to NKp30, NKG2D and CD244 overexpression is not specific to HIV-1 Q23 infection, as we also observed similar responses to mock-infected targets. When co-cultured with mock-infected CD4^+^ T cells, we observed an elevated percentage of functional^+^ NK cells when they overexpress NKp30 or NKG2D (Figures 5D middle panel, S6B and C, bottom panels) and reduced percentage of functional^+^ NK cells when they overexpress CD244 (Figures 5D middle panel, S6D bottom panel). Notably, only 0-5% of NK cells were functional^+^ in both GFP^+^ and GFP^-^ populations when co-cultured with resting CD4^+^ T cells (freshly isolated from PBMCs without stimulation) (Figure 5D bottom panel), indicating that this enhanced activity required B7-H6, MICA/B or ULBP1/2/5/6 expression, which occurs on activated, but not resting, CD4^+^ T cells. Importantly, upregulation of B7-H6 and ULBP1/2/5/6 in infected CD4^+^ T cells is conserved among Q23 and all 4 T/F strains (Figure S2C), suggesting this recognition mechanism is shared across multiple HIV strains.

**Figure 5:**
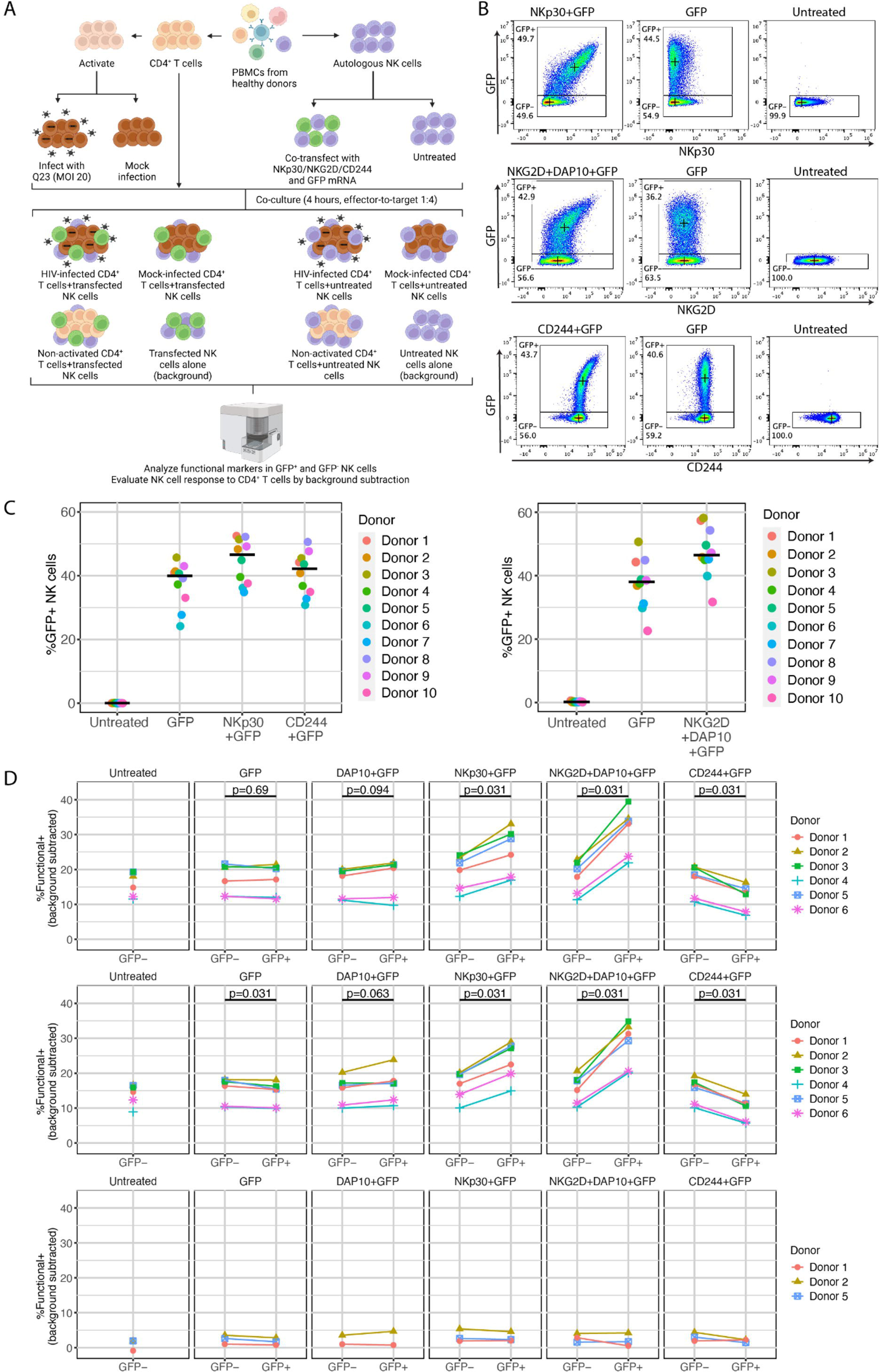
Overexpression of NKp30 or NKG2D enhances NK cell functional response to HIV-infected CD4^+^ T cells. **(A)** A schematic depicting the experimental design of testing the function of transfected NK cells in response to autologous HIV-infected CD4^+^ T cells and mock-infected CD4^+^ T cells. This image was created with BioRender.com. **(B)** Representative flow cytometry plots indicating the co-expression of GFP and each of the (co-)transfected NK cell receptors (NKp30, NKG2D and CD244). Crosses indicate the median fluorescence intensity of the marker on x and y axes in each gate. **(C)** Quantification of the percentage of GFP^+^ NK cells as in (B). n=10. Bars indicate the median of each group. **(D)** Percentage of functional^+^ NK cells (positive for any of CD107a, IFN-D, and TNF-D) in GFP^+^ and GFP^-^ populations after transfection with the mRNA as labeled and co-cultured with autologous Q23-infected (top panel), mock-infected (middle panel) and non-activated CD4^+^ T cells (bottom panel) after subtracting the percentage of functional^+^ NK cells in NK alone group (background subtraction). n=2-6. Statistical analysis was performed with Wilcoxon signed-rank test (nonparametric paired t-test).

Together, these data suggest that NKp30:B7-H6, NKG2D:MICB, and NKG2D:ULBP2 receptor-ligand pairs facilitate NK cell recognition of HIV.

## DISCUSSION

We use mass cytometry to profile in-depth the NK cell response to autologous HIV-infected cells; this encompasses both the phenotypic features of responding NK cells as well as the modulation of cognate NK cell ligand expression on infected CD4^+^ T target cells. To identify the NK cell receptors most important for the response to HIV-infected cells, we further used mass cytometry to characterize both NK cell receptor and ligand expression during NK-CD4^+^ T cell co-cultures. We find that HIV infection induces upregulation of a number of surface proteins that are ligands for activating NK cell receptors, including Nectin-2, B7-H6, CD48, LFA-3 and the various ligands of the NKG2D receptor, though there is considerable variation between donors in the levels of these ligands. Of these, we observed reduced functional NK cell responses to infected cells when eliminating the interactions between NKp30 and its ligand B7-H6, and eliminating the interaction between NKG2D with its ligands MICB or ULBP2 in some donors. Importantly, no single knockout was able to fully abrogate the NK cell response to HIV-infected cells, suggesting redundancy in recognition mechanisms. Consistent with this observation, increasing the interaction between NKp30 and NKG2D and their ligands by overexpressing NKp30 and NKG2D increased the functional NK cell response to HIV-infected cells.

We first optimized an *in vitro* co-culture system to detect NK cell response to HIV-infected cells. It has been previously reported that, *in vitro*, HIV-infected CD4^+^ T cells escape NK cell killing via a number of viral escape pathways mediated by accessory proteins.^15,42^ However, *in vivo*, epidemiological evidence suggests that NK cells can indeed contribute to viral clearance and mediate protection from infection and disease progression, suggesting that *in vitro* methods may require optimization for the effective detection of HIV-specific responses. The use of IL-2 to pre-activate NK cells is known to enhance their ability to lyse infected cells^43^; as such, we used IL-2 activated NK cells to maximize the ability of NK cells to target infected cells, thereby allowing us to probe the specificities of these interactions. While we detected a response to activated, mock-infected autologous CD4^+^ T cells, we also consistently observed a greater response to HIV-infected cells, suggesting that HIV-1 infection drove additional ligand modulation pathways that allowed further targeting by NK cells. Using mass cytometry to compare ligand expression between infected cells, bystander cells and mock-infected cells, we were able to further identify candidate ligands that contributed to recognition of HIV-1 by NK cells, and among these, we demonstrate that overall patterns of B7-H6 and ULBP1/2/5/6 expression were similar across multiple strains of HIV of different subtypes, though there was considerable variation between donors in their expression levels.

We did not account for the effects of KIR and HLA genotypes in our assay, as our primary focus was on identifying shared pathways that were independent of donor genotypes, and hence more broadly applicable in the development of vaccines and therapeutics. Numerous groups have reported the contribution of variation in KIRs and HLAs to both NK cell education and resultant functionality^18^, as well as HIV-targeting NK cell activity. As such, different KIR and HLA haplotypes may contribute to the high degree of variation in NK cell responses observed between donors (Figure 1). However, in our CyTOF screens for receptors that mark HIV-responsive NK cells, the only KIR that was a predictor of functional NK cells was KIR3DL1 (Figure 2A), and it was not a predictor of HIV specificity (vs mock-infected) (Figure 2C). This is consistent with previous reports that KIR3DL1^+^ cells are more functional but are not capable of targeting HIV.^12,44^

Lucar et al reported upregulation of the NKp30 ligand B7-H6 for HIV-2; the authors also showed an upregulation of a NKp30 ligand in HIV-1 infected cells, though to a lower extent.^45^ We observed a small upregulation of B7-H6 in p24^+^ cells compared to mock-infected (Figure 3D), which was supported by the fact that NKp30 was also a predictor of cells responding to HIV-infected cells. In addition, knocking out B7-H6 by CRISPR partially abrogated HIV-specific NK cell response in some donors (Figure S3C).

Overexpression of NKp30 in NK cells elevated NK cell response to Q23-infected CD4^+^ T cells (Figure 5D). These data suggest that B7-H6 can indeed contribute to recognition of HIV-1, similar to HIV-2. However, our ligand panel is limited to the detection of a subset of known ligands for each receptor, and it is unclear if additional NKp30 ligands besides B7-H6, such as BAT3 and galectin-3^46^, may also contribute to recognition of HIV-1. This will be an important area of further investigation.

Among the various NKG2D ligands, ULBPs, but not MICA/B, have been previously reported to be upregulated in HIV-1-infected cells and can trigger NK cell-mediated lysis.^23,47,48^ Here, we confirm that expression of ULBP1/2/5/6 is higher in p24^+^ cells than p24^-^; additionally, we find that the expression of MICA/B is also increased (Figure 3C). The upregulation of MICA/B is possibly specific to certain HIV strains or subtypes since infection with Q23, but not several subtype B strains, either tested in this work or reported previously^47^, led to upregulation of MICA/B in primary CD4^+^ T cells. Antibody blockade of NKG2D was previously shown to reduce NK cell targeting of HIV-infected cells but does not completely abrogate it^23,47,48^; similarly, we find that knockout of individual NKG2D ligands (MICB, ULBP2) in CD4^+^ T cells from most of the donors that were tested induces an incomplete reduction of HIV-specific NK cell responses (Figure S3C) suggesting redundancy, and that all of these ligands contribute to HIV infection recognition by NK cells.

NKp30 and NKG2D overexpression by mRNA and CART transfection enhanced the functional response of NK cells to HIV-infected as well as mock-infected CD4^+^ T cells. Ligands of NKp30 and NKG2D are naturally expressed in tumor, virus-infected and stressed cells, but rarely expressed in normal cells. *In vitro* activation of CD4^+^ T cells can induce the expression of various NKG2D ligands and render CD4^+^ T cells susceptible to NKG2D-mediated NK cell killing.^49^ Here, we activated CD4^+^ T cells with plate-bound α-human CD3 antibodies, PHA-L, and α-human CD28/CD49d antibodies, which renders primary CD4^+^ T cells susceptible to HIV infection *in vitro*. Activated yet uninfected (mock-infected) CD4^+^ T cells also express B7-H6, MICA/B, and ULBP1/2/5/6, although to a lower level than p24^+^ cells (Figure 3D), and interact with NKp30 and NKG2D, which possibly explains the elevation of the function of NKp30 and NKG2D-overexpressing NK cells when co-cultured with mock-infected CD4^+^ T cells. Indeed, when we co-cultured the NK cells that overexpress NKp30 or NKG2D with freshly-isolated non-activated CD4^+^ T cells *in vitro*, NK cell function only marginally increased compared to NK cell alone group, probably because natural CD4^+^ T cells in PBMCs of healthy donors do not express the ligands. Thus, the enhanced responses to mock-infected cells is likely an artifact of the activation necessary to encourage HIV infection.

Receptors on the surface of NK cells, including NKG2D^50–52^, NKp30^53^, and DNAM-1 can be downregulated after persistent binding to their ligands.^54,55^ As such, a limitation of our phenotypic characterization of functional^+^ NK cells (Figure 2) is that we cannot distinguish receptors that are lowly or not expressed, against receptors that were initially expressed on functional^+^ NK cells but were downregulated after ligand engagement. Indeed, we observed reduction of NKG2D expression on the surface of NK cells after they were co-cultured with either HIV-infected or mock-infected CD4^+^ T cells, compared to NK cells that were cultured alone or co-cultured with freshly isolated resting CD4^+^ T cells. In this case, we confirmed the role of NKG2D in facilitating NK cell response to HIV-infected CD4^+^ T cells by co-transfecting NK cells with mRNAs that encode NKG2D and GFP, where GFP consistently indicates the population in which NKG2D level was initially elevated due to transfection, even though NKG2D can be downregulated after co-culturing with HIV-infected or mock-infected CD4^+^ T cells.

CD48 is upregulated after CD4^+^ T cells are infected with the Q23 strain (Figure 3D). When CD244 was overexpressed in NK cells, we observed reduction in the functional response to both Q23-infected and mock-infected CD4^+^ T cells (Figure 5D). Notably, when NK cells transfected with CD244 mRNA were stratified into 3 populations based on the expression level of CD244 (Figure S6A), where CD244^high^ population only includes the cells whose CD244 expression level is higher than any NK cell before transfection, the percentage of functional^+^ cells in CD244^high^ population is significantly higher than in CD244^int^ and CD244^low/-^ populations, while remain similar between CD244^int^ and CD244^low/-^ populations (Figure S6D), suggesting that a hyper physiological level of CD244 inhibits NK cell function. Ezinne et al reported a similar inhibitory role of CD244, where the function of HTLV-1-specific CD8^+^ T cells from both asymptomatic carriers and adult T-cell leukemia/lymphoma (ATLL) patients were elevated with antibody blockade of CD244.^56^ Interaction between CD244 and CD48 can initiate either activating or inhibitory signaling, and the general outcome is determined by the availability of signaling lymphocyte activation molecule (SLAM)-associated protein (SAP). SAP binds to immunotyrosine-based switch motifs (ITSM) on CD244 molecules and transduces activating signals, while in the absence of SAP, SHP-1,2, SHIP-1 and Csk are recruited to ITSM to transduce inhibitory signals.^57–59^ Therefore, when CD48 is upregulated in Q23-infected CD4^+^ T cells, and CD244 is overexpressed in NK cells, that collectively enhances CD244-CD48 interaction, the amount of endogenous SAP may not be sufficient to allow the transduction of activating signals. However, it is possible that the regulation of CD48 is different between HIV strains given the difference we observed with 4 T/F subtype B strains (Figure S2C). Studies on other HIV strains have led to different conclusions about how the level of CD48 expression is influenced during the viral infection.^26,47,60^ Interestingly, Ward et al reported that when CD48 on CD4^+^ T cells is downregulated after infection with two laboratory-adapted macrophage-tropic HIV strains, HIV-1_SF162_ and HIV-1_SF128A_, NK cell killing of HIV-infected T cells is compromised with blockade of CD244^47^, suggesting that low level of CD244-CD48 interaction benefit NK cell activation with the natural amount of SAP. Collectively, the regulation of CD48 in CD4^+^ T cells and the role of CD244-CD48 interaction in NK cell response is possibly specific to HIV strains and warrants further investigation.

A limitation of our study is that we were not able to confirm that knockout of any one NK cell ligand does not impact the expression of others during HIV infection, where accessory protein targeting may be altered in the absence of specific ligands. As such, further studies profiling the entire NK ligand repertoire are required in these edited T cells to better understand potential compensatory mechanisms that may contribute to the limited effects of ligand knockout on NK cell responses we observed (Figure 4C). In addition, knockout of multiple ligands at the same time could delineate the effects of receptor-ligand interactions whose pathways are shared or redundant. In particular, MICA, MICB, ULBP1, and ULBP2 can all induce signaling through the NKG2D receptor, and this redundancy may dampen the effects of knockout of any single ligand.^61^ In addition, our screening of NK cell ligands on infected cells did not account for recognition mechanisms based on potential changes in the peptide repertoire presented on HLA class I molecules in the context of infection. Recent work in non-human primates, as well as humans, have identified SIV/HIV infection-induced alterations in peptides presented on MHC class I^62,63^, suggesting that this may also be an important determinant of NK cell triggering.

The broad genetic variability across HIV strains hinders the development of effective adaptive immune responses.^64,65^ As an innate cell type, NK cells are classically thought to recognize conserved features of pathogens; however, we have previously shown that two strains of influenza can be differentially recognized by NK cells due to differences in ligand modulation.^29^ HIV accessory proteins, many of which are responsible for the modulation of NK ligands, have considerable diversity at the amino acid level^66^, which may lead to differences in ligand modulation between strains. However, using PCA as well as unsupervised clustering, we found that overall patterns of NK ligand expression were determined more by donor (implicating host genetics) than by infecting strain (Figures S2B and 3E). This suggests that methods to target HIV-infected cells via NK recognition mechanisms would be broadly applicable across a variety of strains. Differences in individual ligands have been reported; for example, the commonly used lab-adapted strain NL4-3 does not downregulate HLA-C expression, contrary to other clinical isolate strains.^14^ Indeed, we also observed NL4-3-specific patterns in expression of NK cell ligands - for example, CD48 expression was distinctly higher on NL4-3-infected cells compared to those infected with other viral strains in the same donor (Figure S2C). Overall, methods to target HIV via NK cell receptors are likely to be broadly applicable across HIV strains of different subtypes; however, lab-adapted HIV strains may not entirely recapitulate relevant features in studies of NK cell targeting.

In conclusion, we have developed an experimental and analytical system to probe receptor-ligand interactions involved in the recognition of autologous HIV-infected CD4^+^ T cells by NK cells. By simultaneously profiling NK cell receptor expression and functional activity, and combined with NK ligand expression on infected cells, we were able to develop a list of putative targets that were then investigated by CRISPR knockout of the ligands and overexpression of some of the receptors. This confirmed prior reports of the roles of NKG2D-MICA/B and ULBP1/2/5/6 interactions, and we newly identified NKp30 interaction with B7-H6 as an important mechanism of recognition of HIV-1-infected T cells. While further work is required to delineate the full set of ligands that are required for recognition, our studies set the stage for rationally modulating NK cell ligands and receptors to improve HIV-targeting activity.

## ACKNOWLEDGEMENTS

This work was supported by the National Institutes of Health (NIH DP1 DA046089 and NIH R01 AI161803, C.A.B.), a pilot project through the Bill & Melinda Gates Foundation (OPP1113682, CAB), and the National Science Scholarship from A*STAR Singapore (N.Q.Z.). The Marson lab support includes funding from NIH the HIV Accessory & Regulatory Complexes (HARC) Center (P50 AI150476), NIH/NIAID P01AI138962, Gilead, and The Simons Foundation. We thank Drs. Paul Wender, Gillian Sun and Harrison Rhan in Stanford University for providing the CART ONA used in this work. In addition, we thank the Human Immune Monitoring Core (HIMC) at Stanford University for the use of their Helios mass cytometer.

## Author Contributions

NQZ: Conceptualization; Methodology; Investigation; Formal Analysis; Visualization, Writing - Original Draft

RP: Conceptualization; Methodology; Investigation; Formal Analysis; Visualization, Writing - Original Draft

DNN: Methodology; Investigation; Resources; Writing - Review & Editing

TR: Investigation; Project Administration

CS: Formal Analysis; Software; Writing - Review & Editing

SH: Formal Analysis; Software; Supervision; Writing - Review & Editing

AM: Resources; Supervision; Funding Acquisition; Writing - Review & Editing

CAB: Conceptualization; Supervision; Funding Acquisition; Writing - Review & Editing

## Declaration of Interests

CAB is a scientific advisory board member of ImmuneBridge, DeepCell, Inc., and Qihan Bio on topics unrelated to this manuscript. A.M. is a cofounder of Site Tx, Arsenal Biosciences, Spotlight Therapeutics and Survey Genomics, serves on the boards of directors at Site Tx, Spotlight Therapeutics and Survey Genomics, is a member of the scientific advisory boards of Site Tx, Arsenal Biosciences, Spotlight Therapeutics, Survey Genomics, NewLimit, Amgen, and Tenaya, owns stock in Arsenal Biosciences, Site Tx, Spotlight Therapeutics, NewLimit, Survey Genomics, Tenaya and Lightcast and has received fees from Site Tx, Arsenal Biosciences, Spotlight Therapeutics, NewLimit, 23andMe, PACT Pharma, Juno Therapeutics, Tenaya, Lightcast, Trizell, Vertex, Merck, Amgen, Genentech, GLG, ClearView Healthcare, AlphaSights, Rupert Case Management, Bernstein and ALDA. A.M. is an investor in and informal advisor to Offline Ventures and a client of EPIQ. The Marson laboratory has received research support from the Parker Institute for Cancer Immunotherapy, the Emerson Collective, Juno Therapeutics, Epinomics, Sanofi, GlaxoSmithKline, Gilead and Anthem and reagents from Genscript and Illumina.

**Table S1:**
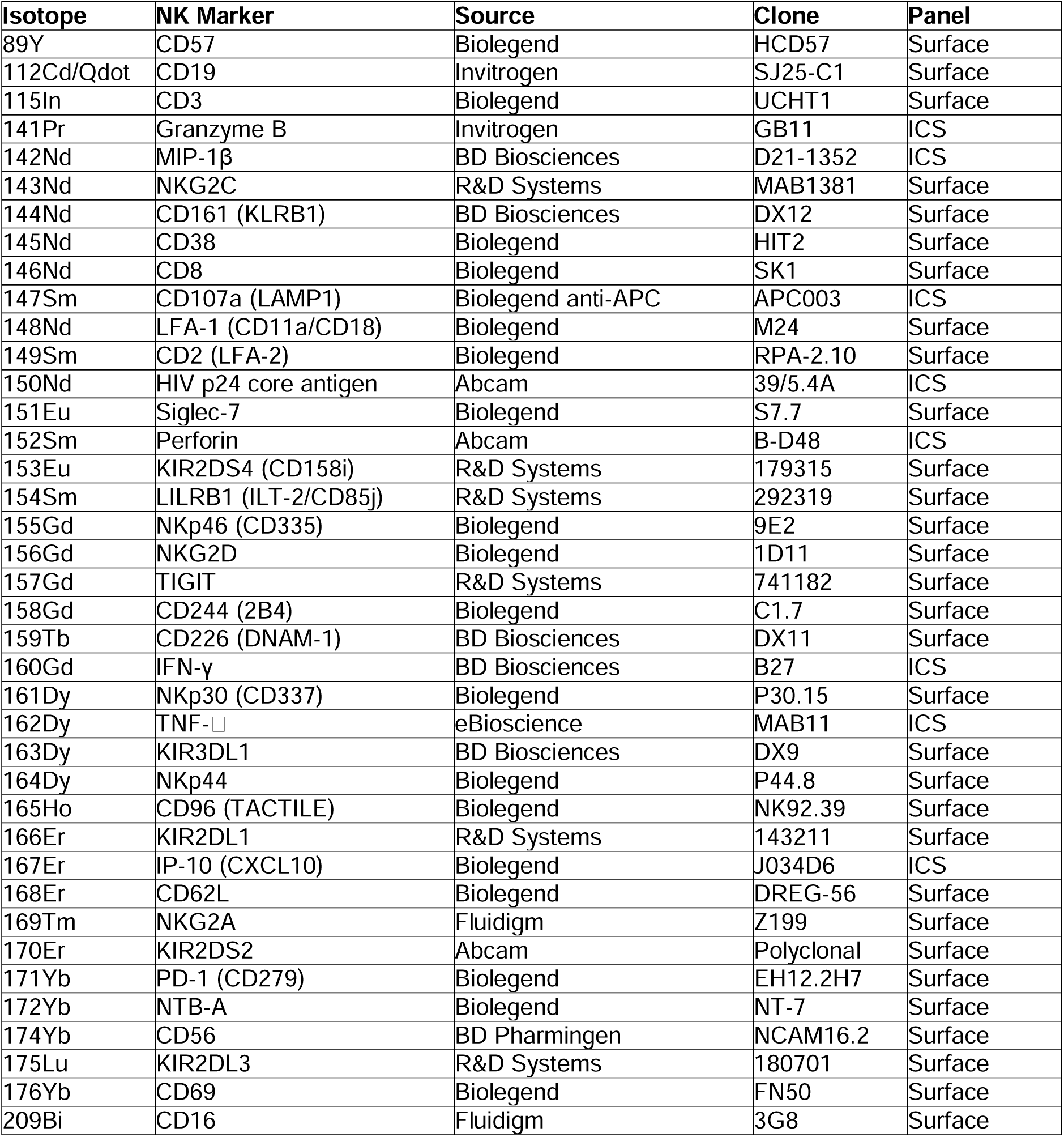
NK CyTOF panel.

| Isotope | NK Marker | Source | Clone | Panel |
| --- | --- | --- | --- | --- |
| 89Y | CD57 | Biolegend | HCD57 | Surface |
| 112Cd/Qdot | CD19 | Invitrogen | SJ25-C1 | Surface |
| 115In | CD3 | Biolegend | UCHT1 | Surface |
| 141Pr | Granzyme B | Invitrogen | GB11 | ICS |
| 142Nd | MIP-1β | BD Biosciences | D21-1352 | ICS |
| 143Nd | NKG2C | R&D Systems | MAB1381 | Surface |
| 144Nd | CD161 (KLRB1) | BD Biosciences | DX12 | Surface |
| 145Nd | CD38 | Biolegend | HIT2 | Surface |
| 146Nd | CD8 | Biolegend | SK1 | Surface |
| 147Sm | CD107a (LAMP1) | Biolegend anti-APC | APC003 | ICS |
| 148Nd | LFA-1 (CD11a/CD18) | Biolegend | M24 | Surface |
| 149Sm | CD2 (LFA-2) | Biolegend | RPA-2.10 | Surface |
| 150Nd | HIV p24 core antigen | Abcam | 39/5.4A | ICS |
| 151Eu | Siglec-7 | Biolegend | S7.7 | Surface |
| 152Sm | Perforin | Abcam | B-D48 | ICS |
| 153Eu | KIR2DS4 (CD158i) | R&D Systems | 179315 | Surface |
| 154Sm | LILRB1 (ILT-2/CD85j) | R&D Systems | 292319 | Surface |
| 155Gd | NKp46 (CD335) | Biolegend | 9E2 | Surface |
| 156Gd | NKG2D | Biolegend | 1D11 | Surface |
| 157Gd | TIGIT | R&D Systems | 741182 | Surface |
| 158Gd | CD244 (2B4) | Biolegend | C1.7 | Surface |
| 159Tb | CD226 (DNAM-1) | BD Biosciences | DX11 | Surface |
| 160Gd | IFN-γ | BD Biosciences | B27 | ICS |
| 161Dy | NKp30 (CD337) | Biolegend | P30.15 | Surface |
| 162Dy | TNF-α | eBioscience | MAB11 | ICS |
| 163Dy | KIR3DL1 | BD Biosciences | DX9 | Surface |
| 164Dy | NKp44 | Biolegend | P44.8 | Surface |
| 165Ho | CD96 (TACTILE) | Biolegend | NK92.39 | Surface |
| 166Er | KIR2DL1 | R&D Systems | 143211 | Surface |
| 167Er | IP-10 (CXCL10) | Biolegend | J034D6 | ICS |
| 168Er | CD62L | Biolegend | DREG-56 | Surface |
| 169Tm | NKG2A | Fluidigm | Z199 | Surface |
| 170Er | KIR2DS2 | Abcam | Polyclonal | Surface |
| 171Yb | PD-1 (CD279) | Biolegend | EH12.2H7 | Surface |
| 172Yb | NTB-A | Biolegend | NT-7 | Surface |
| 174Yb | CD56 | BD Pharmingen | NCAM16.2 | Surface |
| 175Lu | KIR2DL3 | R&D Systems | 180701 | Surface |
| 176Yb | CD69 | Biolegend | FN50 | Surface |
| 209Bi | CD16 | Fluidigm | 3G8 | Surface |

**Table S2:** Ligand CyTOF panel.

| Isotope | NK Ligand | Source | Clone | Panel |
| --- | --- | --- | --- | --- |
| 89Y | HLA-DR | Biolegend | L243 | Surface |
| 115In | CD3 | Biolegend | UCHT1 | Surface |
| 143Nd | pan-HLA class I | Biolegend | W6/32 | Surface |
| 144Nd | CD7 | Biolegend | CD7-6B7 | Surface |
| 145Nd | CD8 | Biolegend | SK1 | Surface |
| 146Nd | CD48 | Biolegend | BJ40 | Surface |
| 148Nd | ICAM1 (CD54) | Biolegend | HA58 | Surface |
| 149Sm | LLT-1 | R&D | 402659 | Surface |
| 150Nd | HIV p24 core antigen | Abcam | 39/5.4A | ICS |
| 151Eu | CD4 | Biolegend | OKT4 | Surface |
| 153Eu | HLA-C | Millipore | DT9 | Surface |
| 154Sm | CCR2 | Biolegend | K036C2 | Surface |
| 155Gd | HLA-E | Biolegend | 3D12 | Surface |
| 156Gd | CD95 | Biolegend | DX2 | Surface |
| 157Gd | Nectin-1 (CD111) | Biolegend | R1.302 | Surface |
| 158Gd | MICA | R&D | 159227 | Surface |
| 158Gd | MICB | R&D | 236511 | Surface |
| 159Tb | DR4 (CD261) | Biolegend | DJR1 | Surface |
| 159Tb | DR5 (CD262) | Biolegend | DJR2-2 | Surface |
| 161Dy | ULBP1 | R&D | 170818 | Surface |
| 161Dy | ULBP2 (5 and 6) | R&D | 165903 | Surface |
| 164Dy | Nectin-2 (CD112) | Biolegend | TX31 | Surface |
| 165Ho | CD155 (PVR) | Biolegend | SKII.4 | Surface |
| 166Er | HLA-Bw4 | Miltenyi | REA274 | Surface |
| 168Er | HLA-Bw6 | Miltenyi | REA143 | Surface |
| 169Tm | CD14 | Biolegend | M5E2 | Surface |
| 170Er | CD11b | Biolegend | ICRF44 | Surface |
| 171Yb | LFA-3 (CD58) | Biolegend | TS2/9 | Surface |
| 172Yb | CD33 | Biolegend | WM53 | Surface |
| 174Yb | CD56 | BD Pharmingen | NCAM16.2 | Surface |
| 176Yb | B7-H6 | R&D | 875001 | Surface |

**Table S3:**
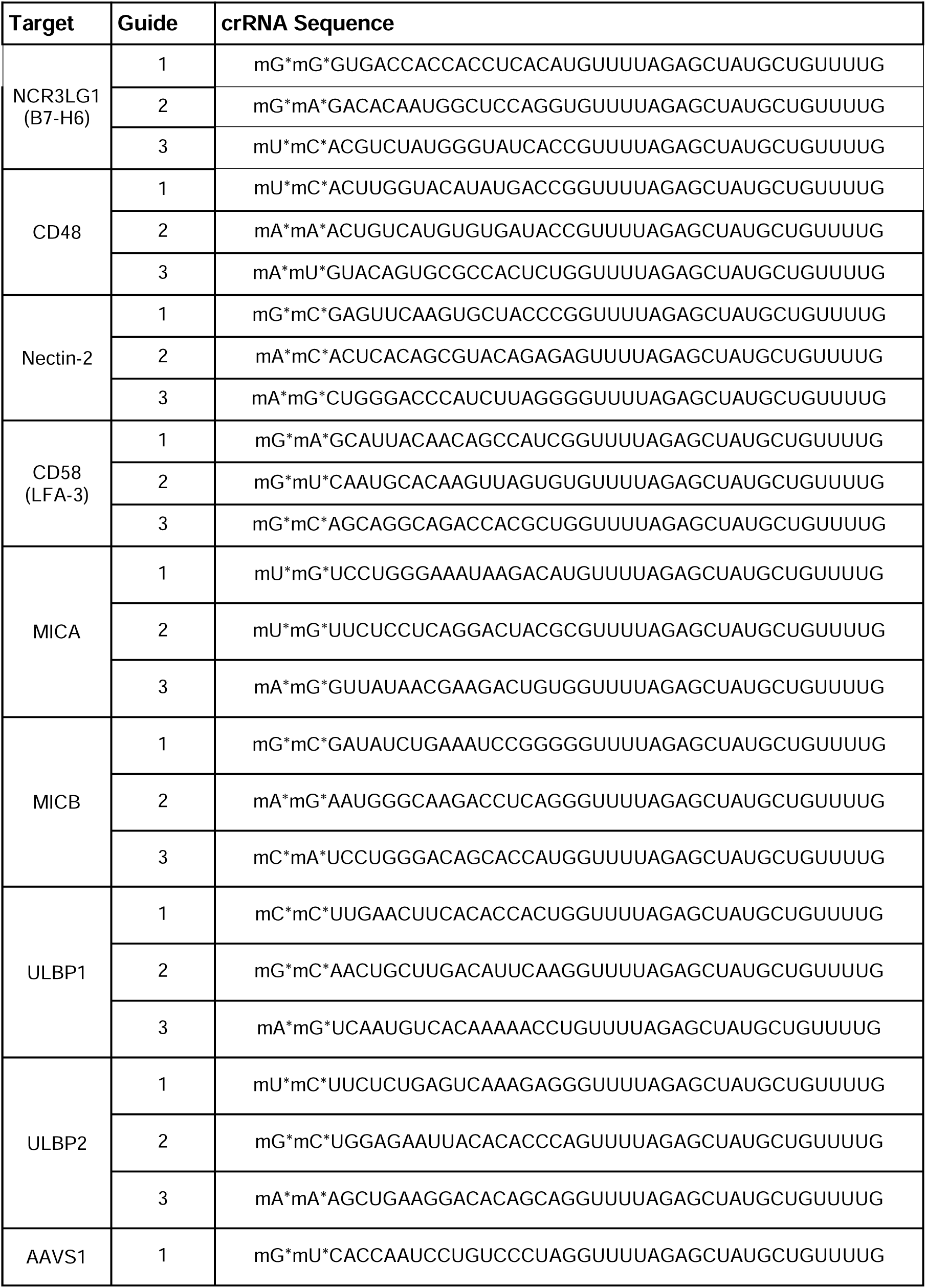
gRNAs used for CRISPR targeting.

## STAR Methods

### KEY RESOURCES TABLE

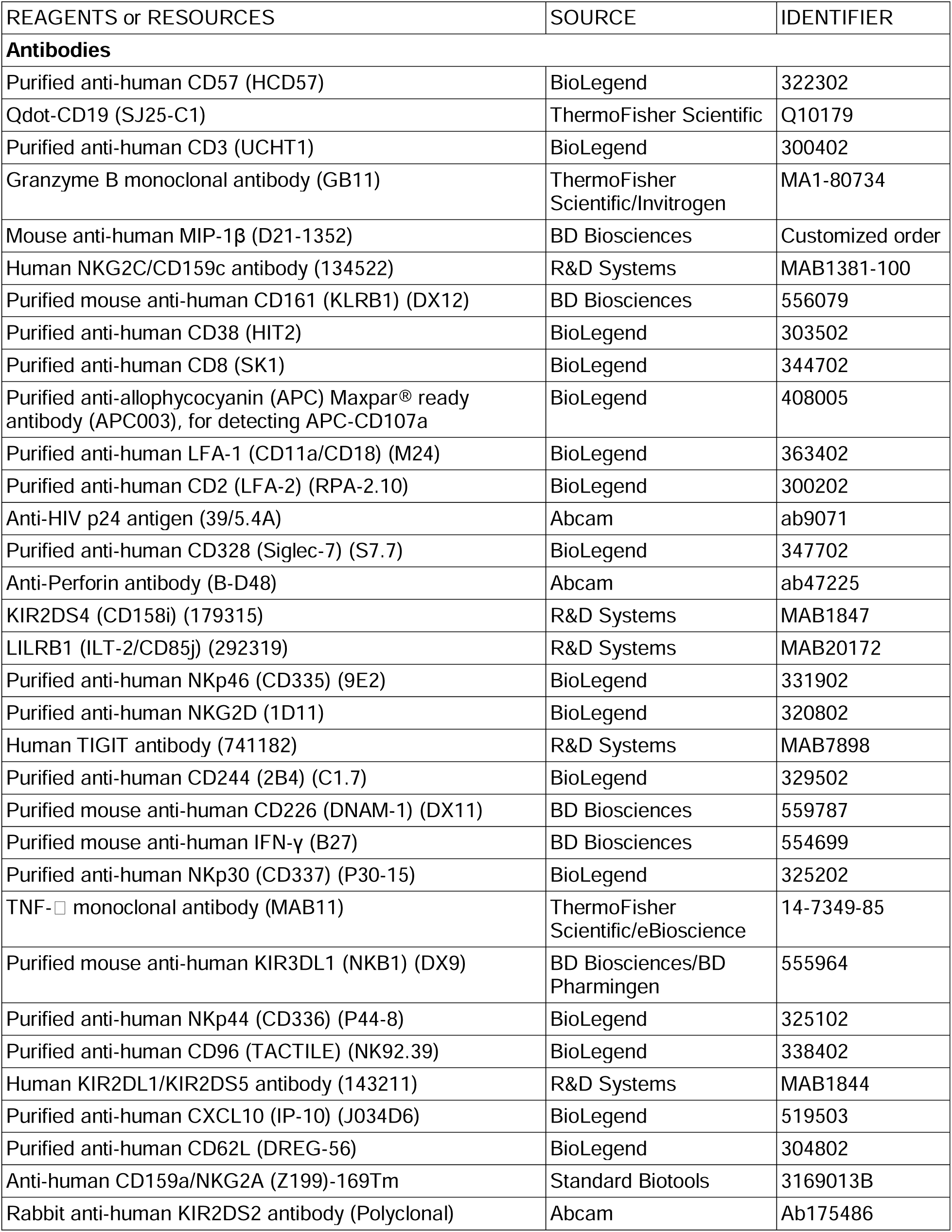
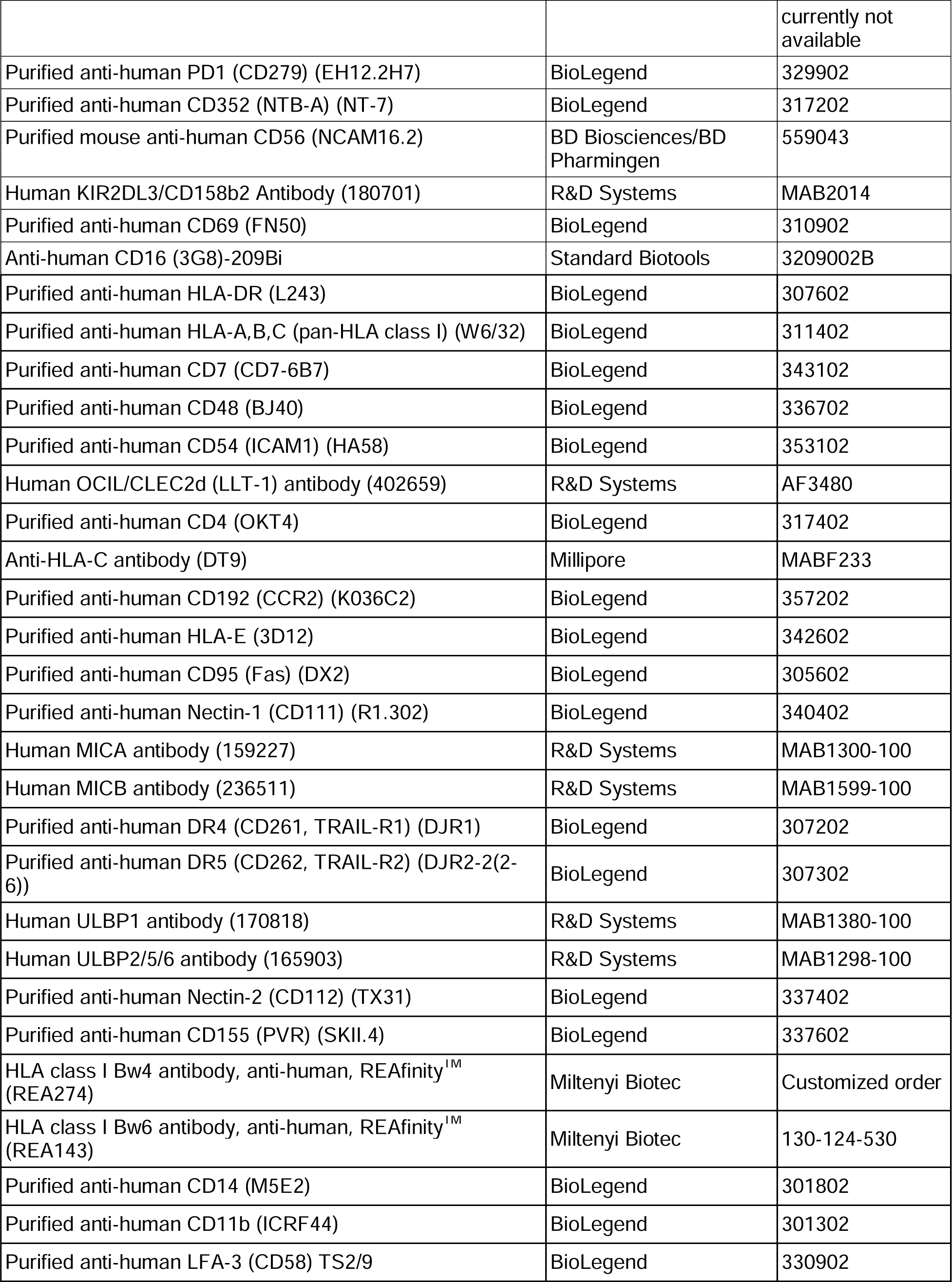
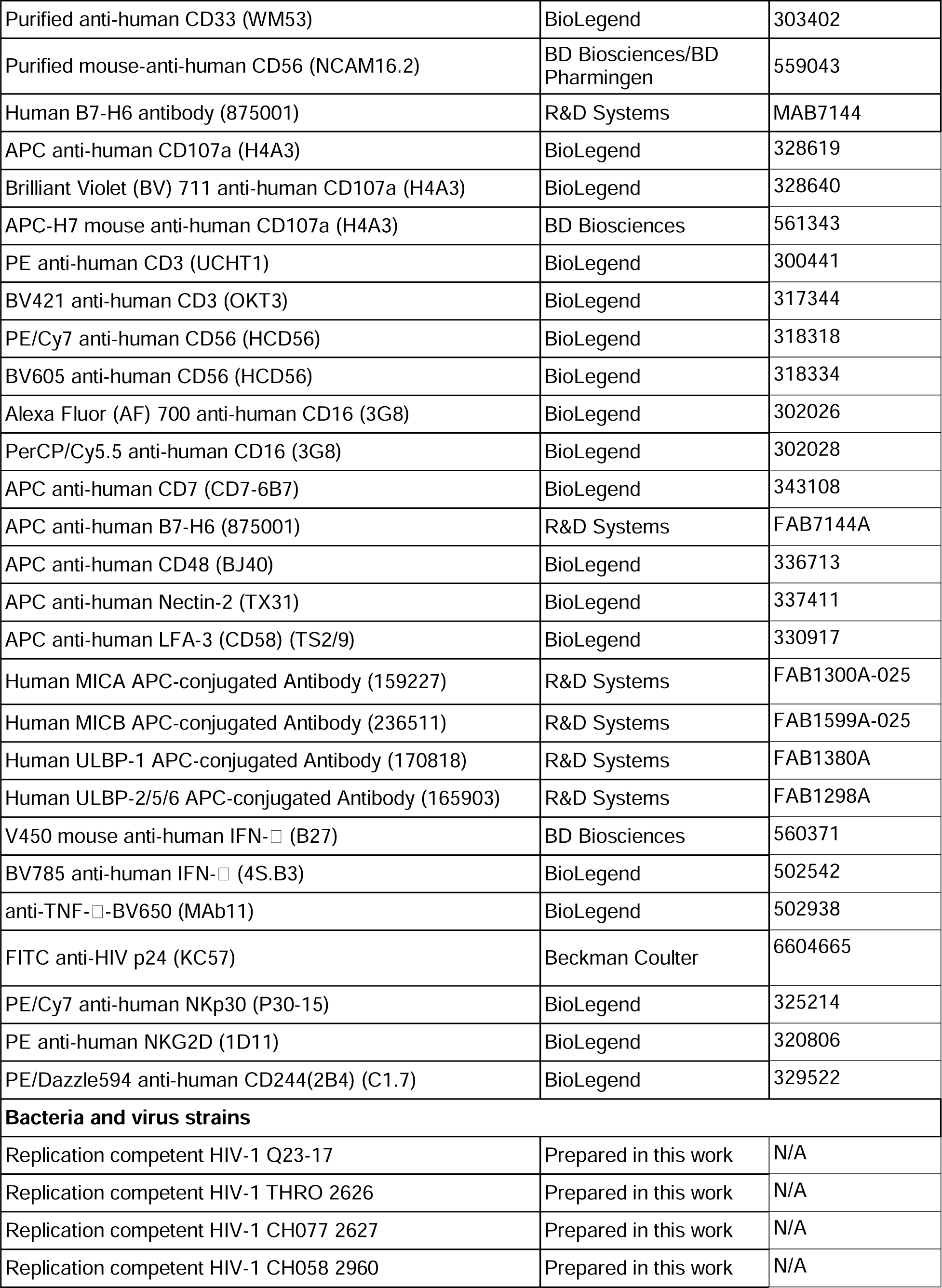
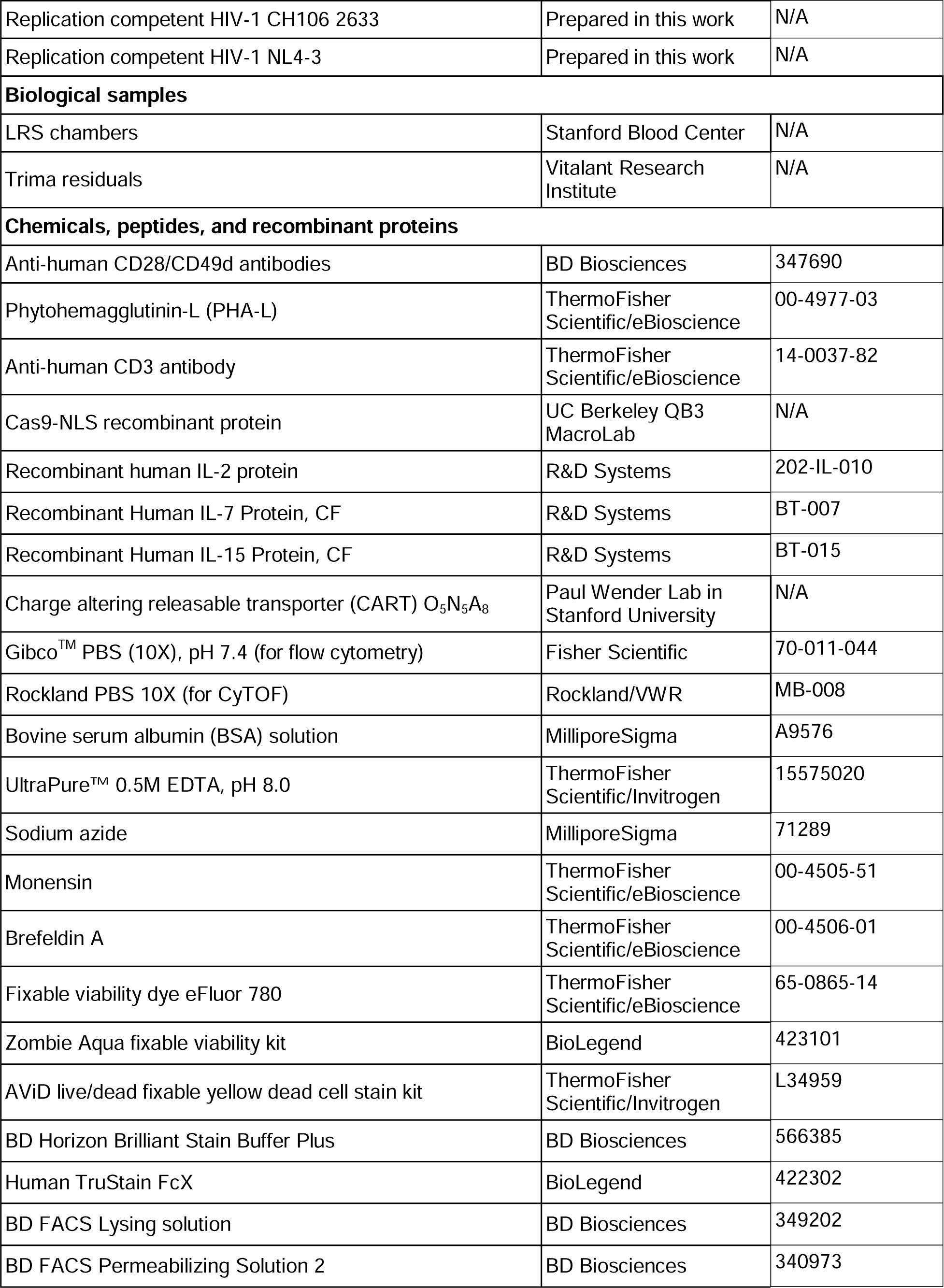
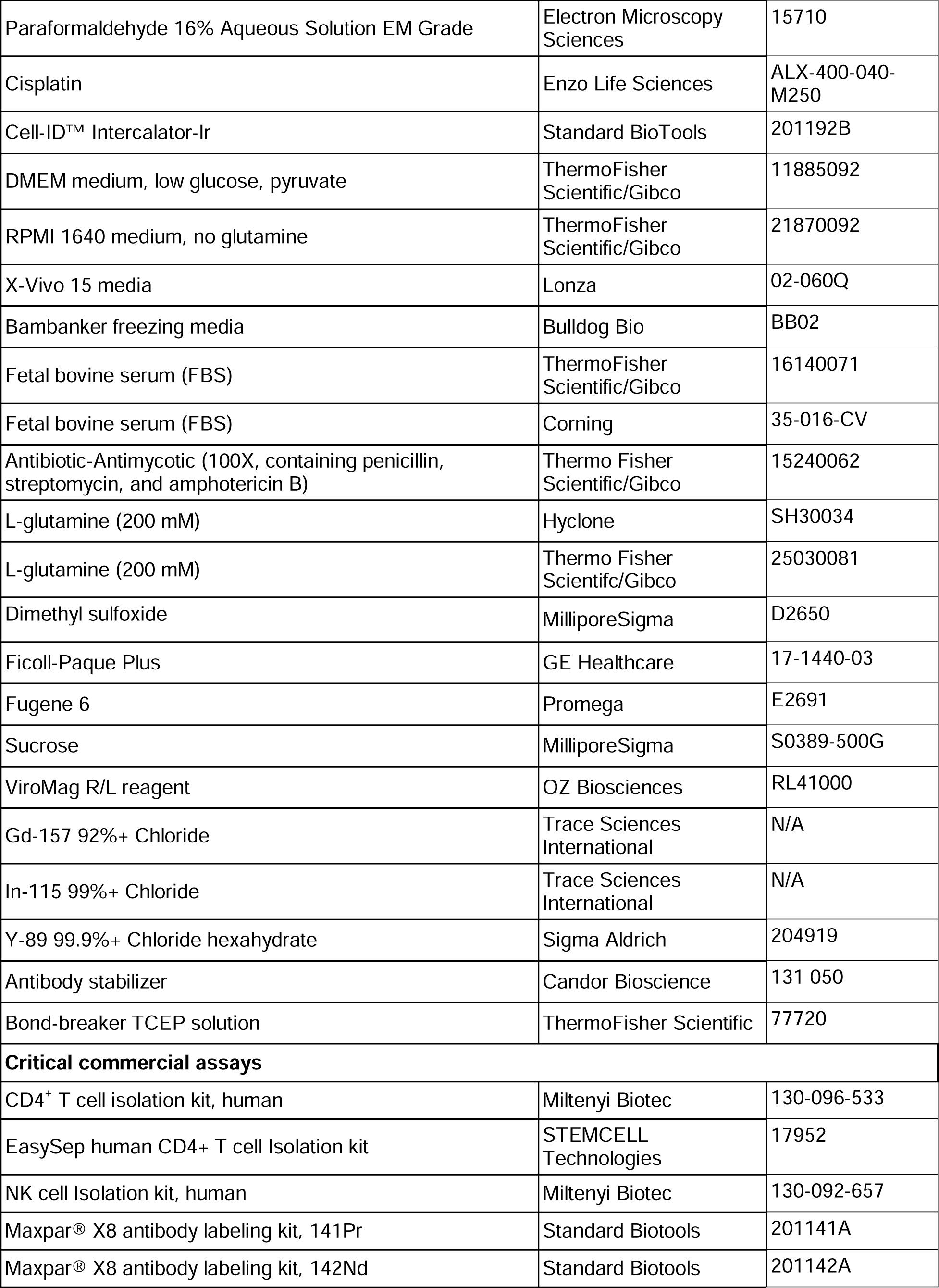
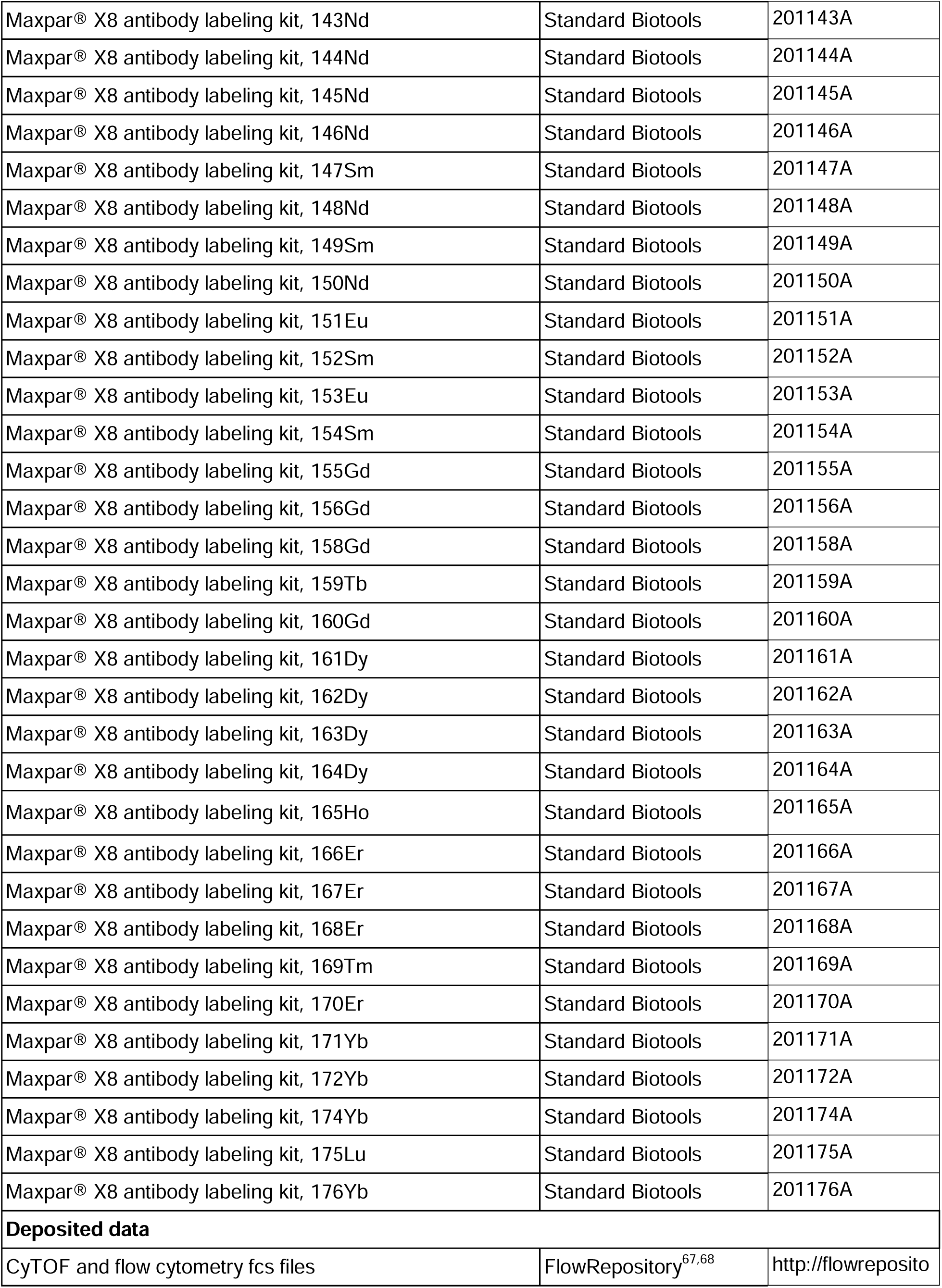
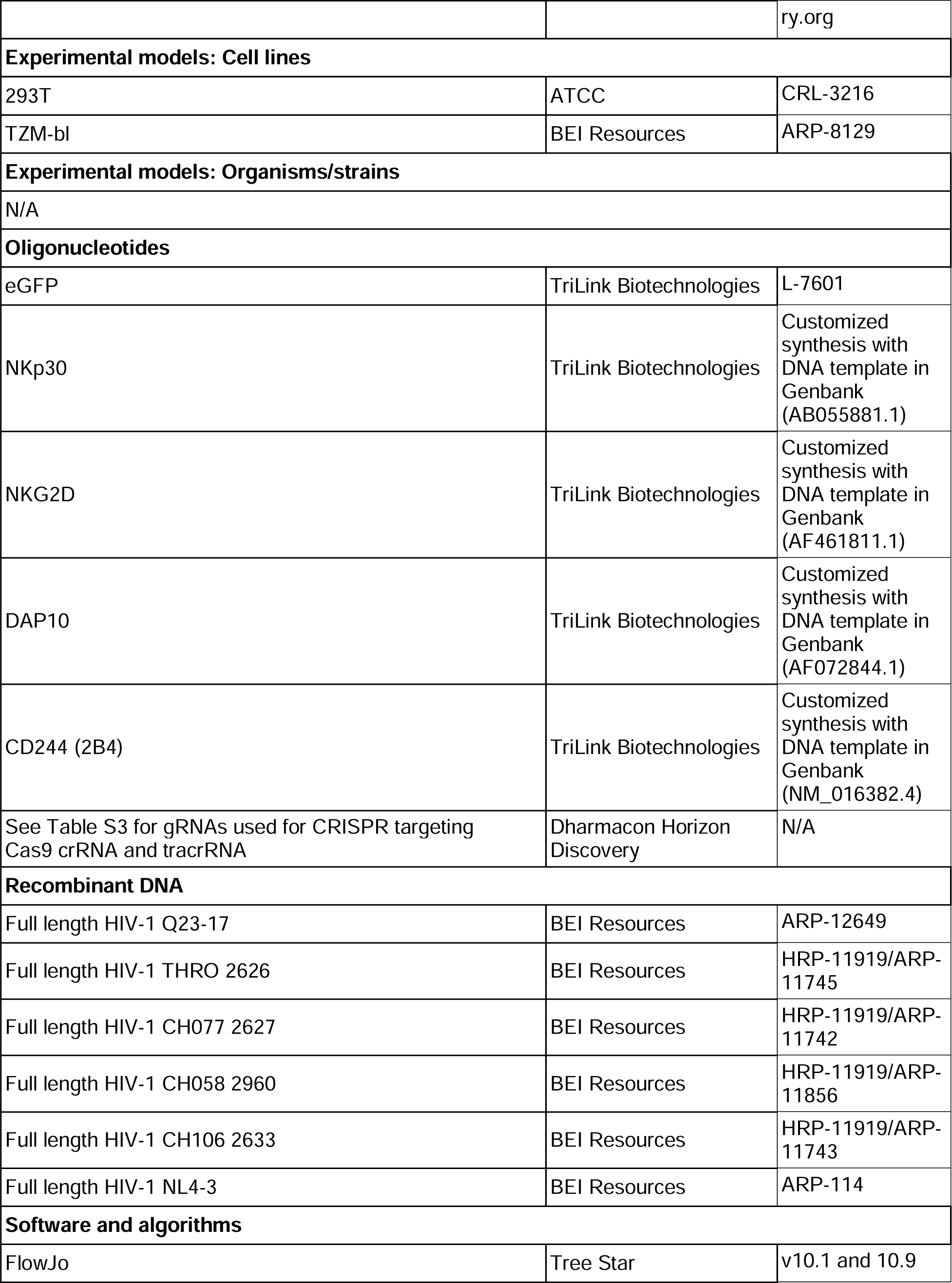
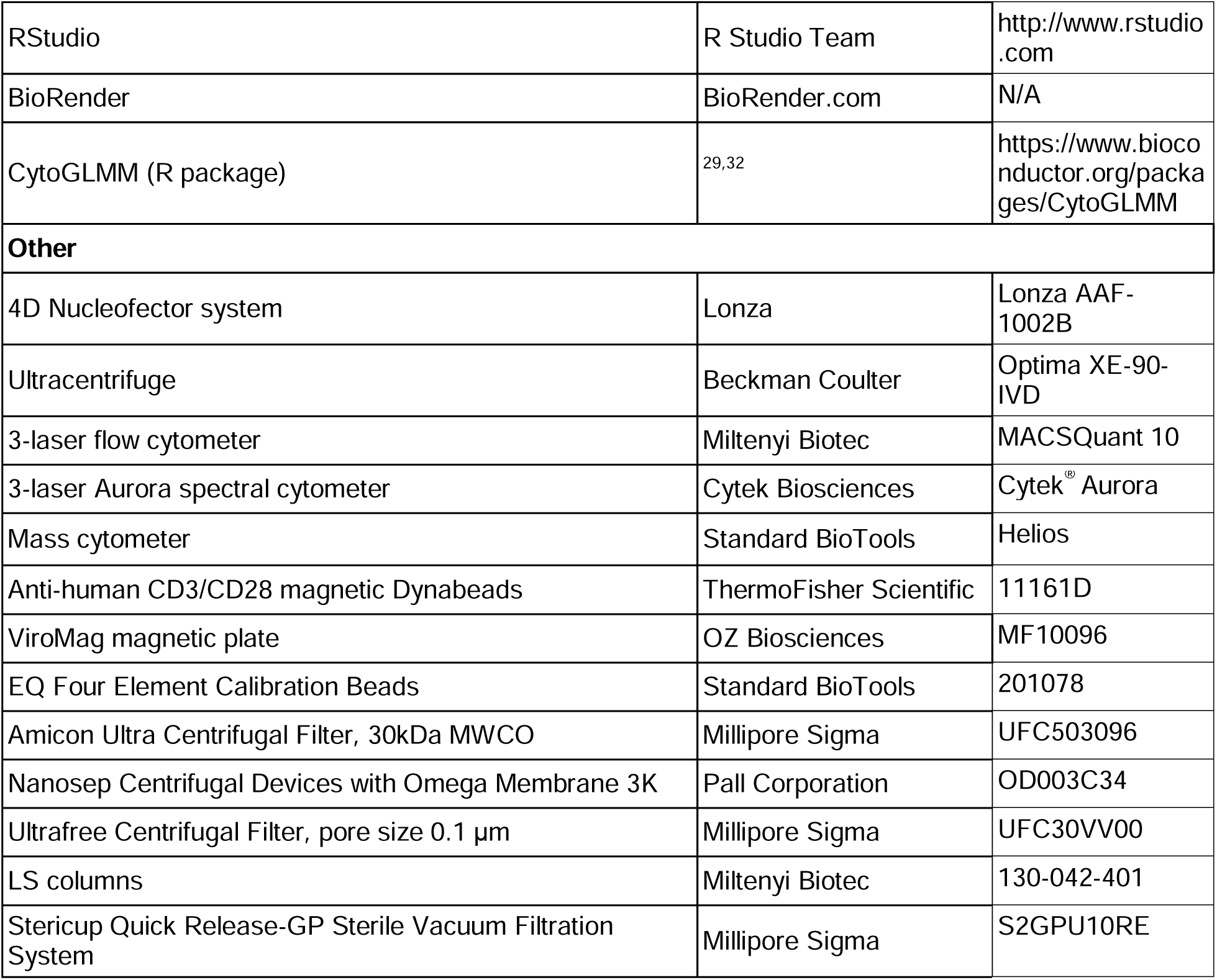

## RESOURCE AVAILABILITY

### Lead contact

Further information and requests for resources and reagents should be directed to and will be fulfilled by the lead contact, Dr. Catherine Blish.

### Materials availability

This study did not generate new unique reagents.

### Data and Code Availability

FCS files of all the CyTOF and flow cytometry analysis in this study can be found in FlowRepository, with the following accession numbers: FR-FCM-Z7H8 for the optimization of CD4^+^ T cells and autologous NK cell *in vitro* co-culture system (Figure 1); FR-FCM-Z7E5 for phenotypic profiling of functional NK cells in response to autologous HIV-infected and mock-infected CD4^+^ T cells (Figures 2, S1A, and S1B); FR-FCM-Z7H9 for the analysis of NK cell ligand expression in CD4^+^ T cells that were infected with Q23 strain (Figures 3A-D and S1C); FR-FCM-Z7G8 for the comparison of NK cell ligand expression in CD4^+^ T cells that were infected with Q23, NL4-3 and a panel of transmitted/founder strains (Figures 3E, S2B and C); FR-FCM-Z7H7 and FR-FCM-Z7H6 for analyzing the expression of NK cell ligands after CRISPR editing of individual genes that encode NK cell ligand (Figures 4B and S3B) and CRISPR editing of AAVS1 (Figure S3A), respectively; FR-FCM-Z7H5 for evaluating the function of NK cell targeting HIV-infected and mock-infected CD4^+^ T cells that were edited with CRISPR/Cas9 (Figures 4C and S3C); FR-FCM-Z7CS for evaluating the expression level of NKp30 and CD244 in NK cells after transfection with CART/mRNA complex (Figures 5B and C); FR-FCM-Z7CU for evaluating the expression level of NKG2D in NK cells after transfection with CART/mRNA complex (Figures 5B, C and S4); FR-FCM-Z7DB for analyzing the function of NK cells that overexpress NKp30, NKG2D, or CD244 in response to HIV-infected, activated mock-infected, and freshly-isolated CD4^+^ T cells (Figures 5D, S5 and S6).^67^

#### EXPERIMENTAL MODEL AND STUDY PARTICIPANT DETAILS

### Cell lines

293T cells were purchased from American Type Culture Collection (ATCC) (catalog number CRL-3216). TZM-bl cells were obtained from BEI Resources (Catalog number ARP-8129). Both 293T and TZM-bl cells were cultured in DMEM media supplemented with 10% FBS (Thermo Fisher) and 1% penicillin/streptomycin/amphotericin (Thermo Fisher).

### PBMC donors

For CRISPR knockout in CD4^+^ T cells and isolation of autologous NK cells, trima residuals were obtained from Vitalant Research Institute (San Francisco, CA) and used for isolating peripheral blood mononuclear cells (PBMCs). For other parts of the study, leukoreduction system (LRS) chambers from healthy, de-identified donors were purchased from Stanford Blood Center and used for isolating PBMCs.

## METHOD DETAILS

### Cell isolation and activation

PBMCs were isolated by density gradient centrifugation using Ficoll-Paque PLUS (GE Healthcare), and cryopreserved in 10% DMSO (Sigma Aldrich) and 90% FBS (Thermo Fisher). For all the assays, PBMCs were thawed, and NK cells and CD4^+^ T cells were separately isolated by negative selection with kits purchased from Miltenyi Biotec unless otherwise indicated. All cells were cultured in RPMI (Gibco), with 10% FBS (Thermo Fisher), 1% L-glutamine (Hyclone) and 1% penicillin/streptomycin/amphotericin (Thermo Fisher) (RP10). NK cells were cultured in RP10 supplemented with 300 IU/ml recombinant human IL-2 (R&D Systems) for 72 h unless otherwise noted. CD4^+^ T cells were activated in RP10 with plate-bound anti-CD3 (clone OKT3, eBioscience, plates coated at 10 μg/ml), anti-CD28/CD49d (BD Biosciences, final concentration 1 μg/ml per antibody) and PHA-L (eBioscience, final concentration 3.1 μg/ml) for 48 h in all the experiments unless otherwise noted.

### HIV preparation, titration and infection in CD4^+^ T cells

Multiple strains of HIV were prepared by transfecting plasmids encoding full-length, replication competent clones into 293T cells with Fugene 6 (Promega). Plasmids encoding full-length Q23-17 (shortened as Q23), a subtype A strain cloned from a patient one year after seroconversion^31^, subtype B transmitted/founder (T/F) strains, including THRO 2626, CH077 2627, CH058 2960 and CH106 2633 as well as subtype B laboratory-adapted strain NL4-3 were obtained from BEI Resources. Supernatant from 293T cultures was harvested 48 h after transfection and viruses were concentrated by ultracentrifugation. Viral stocks were titrated on TZM-bl cells as previously described.^69^ Activated CD4^+^ T cells were infected with HIV using Viromag magnetofection (OZ Biosciences). Q23 was added into CD4^+^ T cells at a multiplicity of infection (MOI) of 20 in all the experiments except for the analysis of CD4^+^ T cells infected with different HIV strains (Figures 3E and S3). To compare how different HIV strains modulate NK ligand expression in CD4^+^ T cells, 6 HIV strains, including Q23, THRO 2626, CH077 2627, CH058 2960, CH106 2633, and NL4-3 were added to CD4^+^ T cells at an MOI of 10, 10, 10, 20, 4, and 0.5, respectively, to achieve a similar level of infection (p24^+^ in 40-80% of total CD4^+^ T cells identified with flow cytometry). CD4^+^ T cells that were treated with the same amount of Viromag reagent in absence of HIV (mock-infection) served as a control group. HIV-infected and mock-infected cells were used for co-culture with NK cells or stained with the ligand panel described in Table S2 for mass cytometry (CyTOF) analysis 24 h post infection.

### CRISPR knockout in CD4^+^ T cells

Trima residuals were obtained from Vitalant (San Francisco, CA) and PBMCs separated using Ficoll. 3x10^8^ PBMCs were cryopreserved in Bambanker freezing media (Bulldog Bio, catalog number BB02) for later NK cell isolation. CD4^+^ T cells were isolated using the EasySep Human CD4^+^ T cell Isolation Kit (STEMCELL Technologies), and activated with anti-CD3/anti-CD28 Dynabeads (Gibco) at a 1:1 bead:cell ratio, in X-Vivo 15 media (Lonza) supplemented with 300 IU/ml IL-2, 5ng/ml IL-7 and 5ng/ml IL-15 for 48 h. Subsequently, activated CD4^+^ T cells were de-beaded, incubated with Cas9 ribonucleoproteins targeting each gene of interest, and electroporated using a 4D Nucleofector system (Lonza) as previously described.^70^ Details of all gRNAs are given in Table S3. Post-electroporation, cells were maintained for 5 days in X-Vivo 15 media supplemented with 500 IU/ml IL-2. Cells were then infected with Viromag magnetofection, as described above, with the Q23 virus.

### NK cell transfection with CART

NK cells were isolated from PBMCs of healthy donors as mentioned above. NK cells were washed twice with serum-free RPMI (Gibco) media (supplemented with 1% L-glutamine and 1% penicillin/streptomycin/amphotericin), resuspended in serum-free RPMI media and plated in a non-tissue culture treated U-bottom 96-well plate at density of 500,000 NK cells/100 μl in each well for mRNA transfection with CART. mRNA fragments that encode NKp30, NKG2D, DAP10, and CD244 were synthesized at TriLink Biotechnologies with DNA templates of coding sequences (cds) obtained from GenBank® database. During the mRNA synthesis, uridine and cytidine were fully substituted with pseudouridine and 5-methylcytidine. A cap 1 structure (CleanCap AG) as well as a 120-nucleotide polyadenosine (polyA) tail were incorporated into the mRNA fragments to reduce the immunogenicity and enhance the stability of the mRNA. GFP mRNA (catalog number L-7601) was purchased from TriLink Biotechnologies. CARTs are polycationic oligomers that can package polyanionic nucleic acids and deliver the nucleic acids into cells. Once in the cells, CARTs are degraded into nontoxic small molecules and release the nucleic acids.^38^ For co-delivering mRNA that encodes GFP with mRNA that encodes NKp30, NKG2D, CD244, or DAP10, stock of mRNA was premixed to include a combination of NKp30 and GFP mRNA (mass ratio 1:1) or a combination of NKG2D, DAP10, and GFP mRNA (mass ratio 3:1:3 or otherwise as indicated) or a combination of 2B4 and GFP mRNA (mass ratio 1:1) or a combination of DAP10 and GFP mRNA (mass ratio 1:6). The CART/mRNA complexes were prepared by mixing a CART named O_5_:N_5_:A_8_ (shortened as ONA) with each of the premixed mRNA combinations or GFP mRNA alone according to the protocol that was optimized previously and was added into each well of NK cells in the 96-well plate at a dosage of 100 ng mRNA per well.^39^ GFP mRNA that was not mixed with ONA was added into NK cells as an untreated control group. Two hours later, media in the 96-well plate was changed to complete RPMI media that contains 10% FBS, 1% L-glutamine and 1% penicillin/streptomycin/amphotericin without adding any cytokine. NK cells were cultured overnight and co-cultured with HIV-infected or mock-infected CD4^+^ T cells.

### CD4^+^ T-NK cell co-culture

NK cells and CD4^+^ T cells were co-cultured for 4 h, in the presence of brefeldin A (eBioscience, final concentration 3.0 μg/ml), monensin (eBioscience, final concentration 2 μM), and anti-CD107a-APC (for CyTOF analysis in Figure 2), or anti-CD107a-APC-H7 (Figures 1 and 4), or anti-CD107a-BV711 (Figure 5D) at effector:target (E:T) ratio as indicated. At the end of the co-culture, cells were analyzed with flow cytometry or CyTOF.

### Flow cytometry analysis

Cells were first stained with a viability dye, then stained with antibodies that bind to molecules on the cell surface. For the analysis of NK cell functional markers and killing of HIV-infected CD4+ T cells in the *in vitro* co-culture system (Figure 1), cells were first stained with AViD Live/Dead Fixable Yellow Dead Cell Staining Reagent and subsequently stained with PE anti-CD3 (Biolegend, clone UCHT1), PE/Cy7 anti-CD56 (Biolegend, clone HCD56), PerCP/Cy5.5 anti-CD16 (Biolegend, clone 3G8), and APC anti-human CD7 (BioLegend, clone CD7-6B7). For the analysis of NK cell ligand expression in CD4^+^ T cells or functional analysis of NK cells co-cultured with autologous CRISPR edited CD4^+^ T cells (Figure 4), cells were first stained with Zombie Aqua Fixable Viability dye (Biolegend) and subsequently stained with PE anti-CD3 (Biolegend, clone UCHT1), PE/Cy7 anti-CD56 (Biolegend, clone HCD56) and APC anti-ligand (B7-H6: R&D, clone 875001; CD48: Biolegend, clone; BJ40; Nectin-2: Biolegend, clone TX31; LFA-3: Biolegend, cloneTS2/9; MICA: R&D, clone 159227; MICB: R&D, clone 236511; ULBP1: R&D, clone 170818; ULBP2/5/6: R&D, clone 165903; all ligand antibodies used the same clones as the CyTOF ligand panel in Table S2). For the analysis of NKp30, NKG2D, and CD244 expression in NK cells after CART transfection (Figures 5B, C, S4), NK cells were first stained with fixable viability dye eFluor 780, then treated with Human TruStain FcX for blocking Fc receptors and stained with Brilliant Violet (BV) 421 anti-CD3 (BioLegend, clone OKT3), APC anti-human CD7 (BioLegend, clone CD7-6B7), BV605 anti-CD56 (BioLegend, clone HCD56), Alexa Fluor (AF)700 anti-CD16 (BioLegend, clone 3G8), PE/Cy7 anti-NKp30 (BioLegend, clone P30-15), PE anti-NKG2D (BioLegend, clone 1D11) and PE/Dazzle594 anti-CD244 (BioLegend, clone C1.7) in the presence of BD Horizon brilliant stain buffer plus (BD Biosciences). For the functional analysis of CART/mRNA-transfected NK cells in the co-culture with CD4^+^ T cells (Figures 5D, S5, and S6), cells were first stained with fixable viability dye eFluor 780 and stained with the same panel of antibodies as for analyzing the expression of NKp30, NKG2D and CD244 that was mentioned above, except for APC anti-human CD7.

For the samples that require intracellular staining, cells were subsequently fixed with FACS Lyse (BD Biosciences), permeabilized with FACS Permeabilization Buffer 2 (BD Biosciences), and stained for intracellular markers with V450 anti-IFN-D (BD Biosciences, clone B27) or BV785 anti-IFN-D, BV650 anti-TNF-D (Biolegend, clone MAb11) and FITC HIV p24 (Beckman Coulter, clone KC57) as indicated. Cells were fixed with 1% or 2% paraformaldehyde (PFA) in PBS and analyzed by flow cytometry using an Aurora spectral cytometer (Cytek Biosciences), and data analysis was performed using FlowJo version 10.1 and 10.9 (Tree Star).

### Mass cytometry (CyTOF) staining and sample acquisition

All CyTOF antibodies were conjugated using MaxPar® X8 labeling kits (Standard BioTools), except for those purchased directly from Standard BioTools; details of all antibodies are given in Table S1 (NK panel) and Table S2 (ligand panel). Table S1 describes a panel of antibodies for analyzing NK cell receptor expression in NK cells after CD4^+^ T-NK cell co-culture (Figure 2). Qdot anti-human CD19, 141Pr anti-human Granzyme B, 142Nd anti-human MIP-1β, 148Nd anti-human LFA-1, 152Sm anti-human Perforin, 167Er anti-human IP-10, and 170Er anti-human KIR2DS2 were not included in data analysis. Table S2 describes a panel of antibodies for analyzing the expression of NK ligands on CD4^+^ T cells after HIV-infection or mock-infection (Figures 3 and S2). 144Nd anti-human CD7, 154Sm anti-human CCR2, 166Er anti-human HLA-Bw4, 168Er anti-human HLA-Bw6, 169Tm anti-human CD14, 172Yb anti-human CD33, and 174Yb anti-human CD56 were not included in data analysis. To maintain antibody stability and consistency in staining, all antibody panels were pre-mixed into separate surface and ICS cocktails (as indicated in the Tables), aliquoted and frozen at -80°C until use. At the end of HIV-infection or CD4^+^ T-NK cell co-culture, cells were stained for viability using 25 μM Cisplatin (Enzo) for 1 min and quenched with FBS, and washed with CyFACS buffer (1X Rockland PBS with 0.1% BSA, 2mM EDTA and 0.05% sodium azide). Samples were stained with the surface antibody panel for 30 min at 4°C, fixed with FACS Lyse (BD Biosciences), permeabilized with FACS Permeabilization Buffer 2 (BD Biosciences), and stained with the intracellular staining (ICS) panel for 45 min at 4°C. Cells were suspended in and treated with iridium intercalator (Standard BioTools) in 2% paraformaldehyde (PFA) overnight, and resuspended in 1x EQ Beads (Standard BioTools) before acquisition on a Helios mass cytometer (Standard BioTools).

## QUANTIFICATION AND STATISTICAL ANALYSIS

### CyTOF data analysis

Bead normalization (https://github.com/nolanlab/bead-normalization) was performed on all files for each set of experiments post-acquisition. All CyTOF data was visualized and gated using FlowJo v10.1 (Tree Star); gated NK cells (CD3^-^ CD56/CD16^+^), bulk CD4^+^ T cells (CD3^+^ CD8^-^; CD4 was not used for gating as CD4 downregulation can occur after HIV infection) or p24^+/-^ CD4^+^ T cells were exported as fcs files from FlowJo and used in downstream analyses (gating schemes in Figure S1). The open source statistical software R was used for analyses.^71^

To visualize the multivariate CyTOF data, we used principal components analysis (PCA) on the per sample median marker expression.^72^ We color coded each sample with infection virus strain and blood donor. We used a circle plot to show the correlations of the PCA axes with each marker. To increase interpretability, we focused on markers that have a correlation coefficient of 0.5 or greater along either PCA axes. Additionally, we used a heatmap on the mean asinh-transformed expression with a dendrogram derived from unsupervised clustering.^72^ To improve the visual comparison between markers, we standardized expressions across samples.

To identify phenotypic markers predictive of functionally responding NK cells, HIV-infected CD4^+^ T cells, or HIV antigen expressing (p24^+^)^-^ CD4^+^ T cells, we used the R package *CytoGLMM* that uses a generalized linear mixed model.^29,32^ This model takes into account the distribution of each marker and has a donor-specific variable to control for inter-individual variability. For analyses using *CytoGLMM*, raw channel values were transformed using the inverse hyperbolic sine (asinh) function with a cofactor of 5 to account for heteroskedasticity. This transformation was not applied for calculating mean signal intensity values.

