## Supplementary figures for "NKp30 and NKG2D contribute to natural killer recognition of HIV-infected cells"

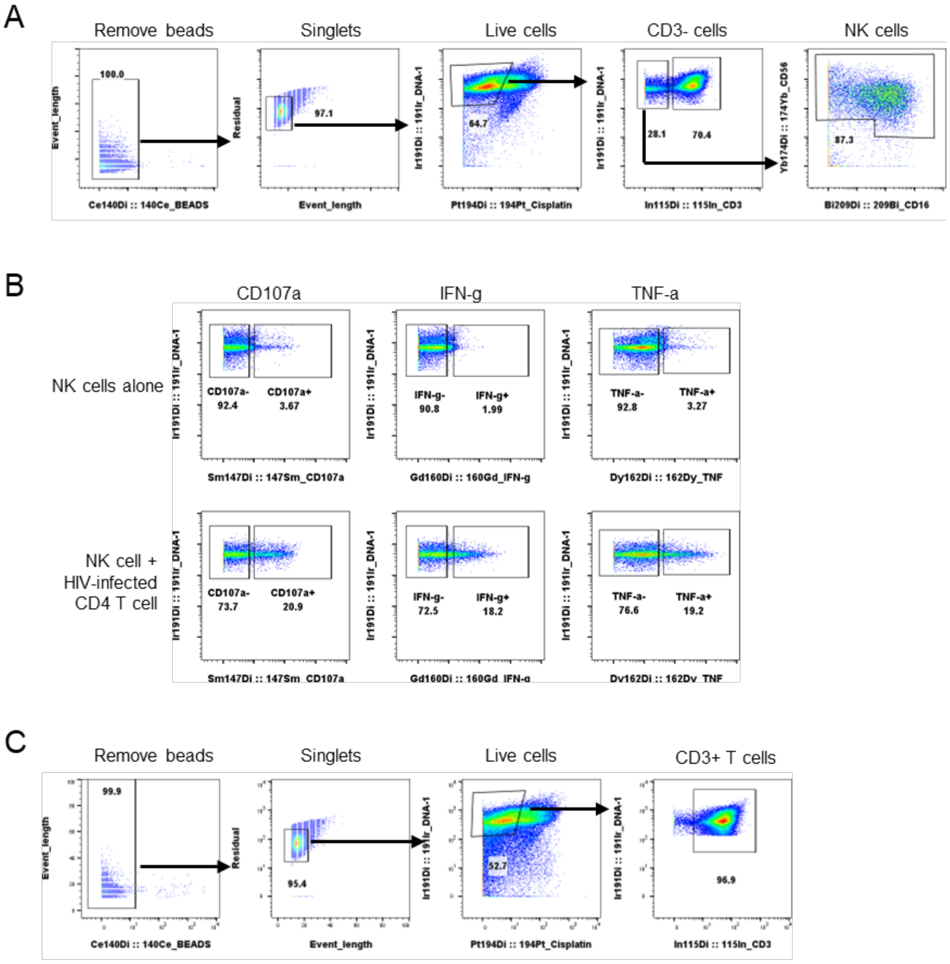


Figure S1

**A.** Gating strategy for NK cells in CyTOF data. **B.** Representative flow cytometry plots of CD107a, IFN-γ, and TNF-α production (by frequency of positive cells), after 4 h co-culture without (top) or in the presence of HIV-infected autologous $CD4^+$ T cells (bottom). **C.** Gating strategy for $CD4^+$ T cells in CyTOF data.

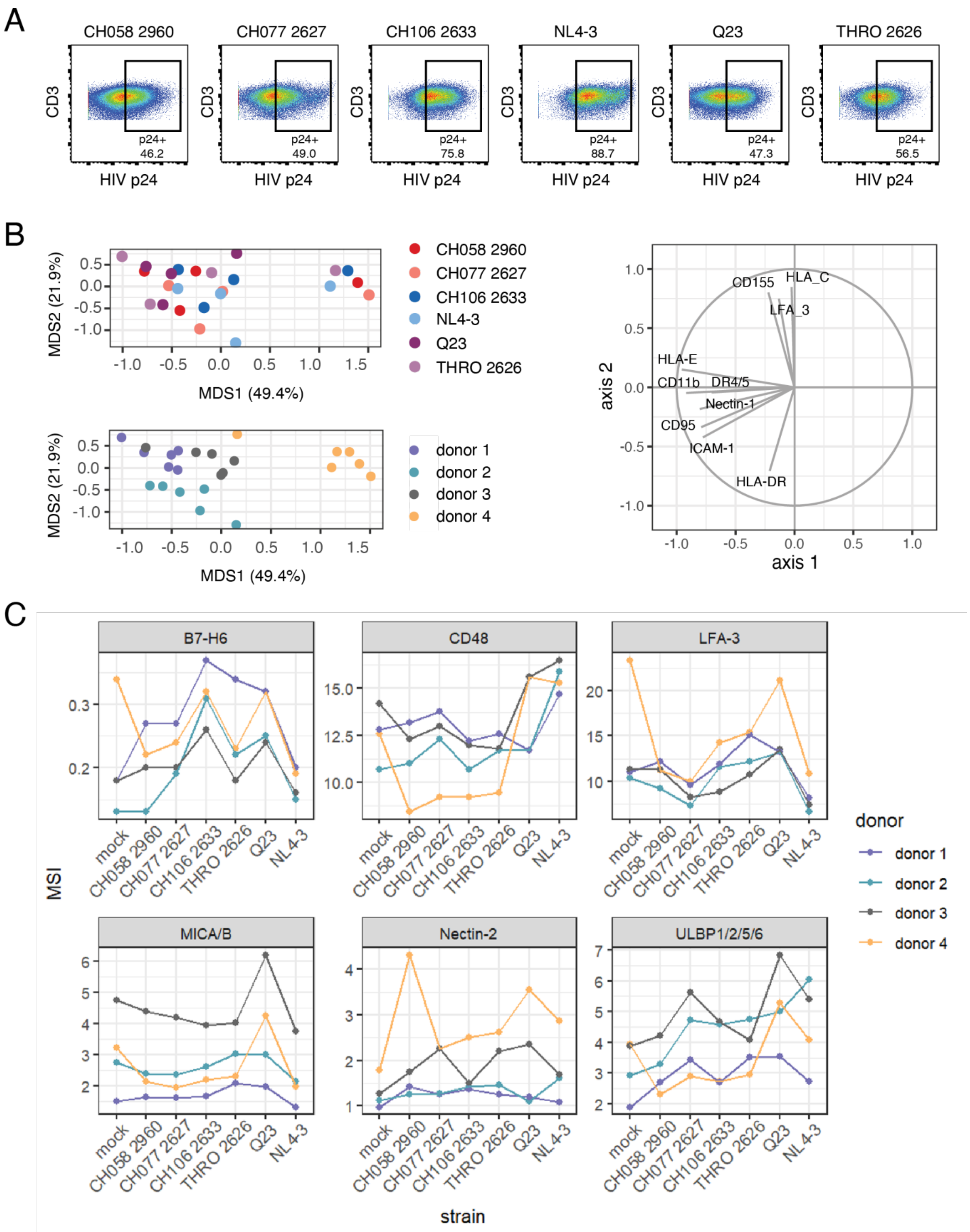


Figure S2. HIV strain and *donor-dependent variability of NK cell ligand modulation in infected CD4⁺ T cells.*

**(A)** Representative dot plots from mass cytometry, demonstrating a similar level of infection of primary CD4$^+$ T cells with all HIV strains tested, as measured by percentage positively staining for HIV p24. **(B)** Principal components analysis (PCA) plot showing separation of all samples, gated to p24$^+$ HIV-infected cells, with each sample coloured by infecting virus strain (top), or blood donor (bottom). Only markers whose contributions are greater than 0.5 in either PCA1 or PCA2 are displayed in the marker loadings. **(C)** Mean signal intensity (MSI) of NK cell ligands on mock-infected CD4$^+$ T cells and p24$^+$ CD4$^+$ T cells infected with each HIV strain as indicated. n=4. Each individual donor is joined with a line.

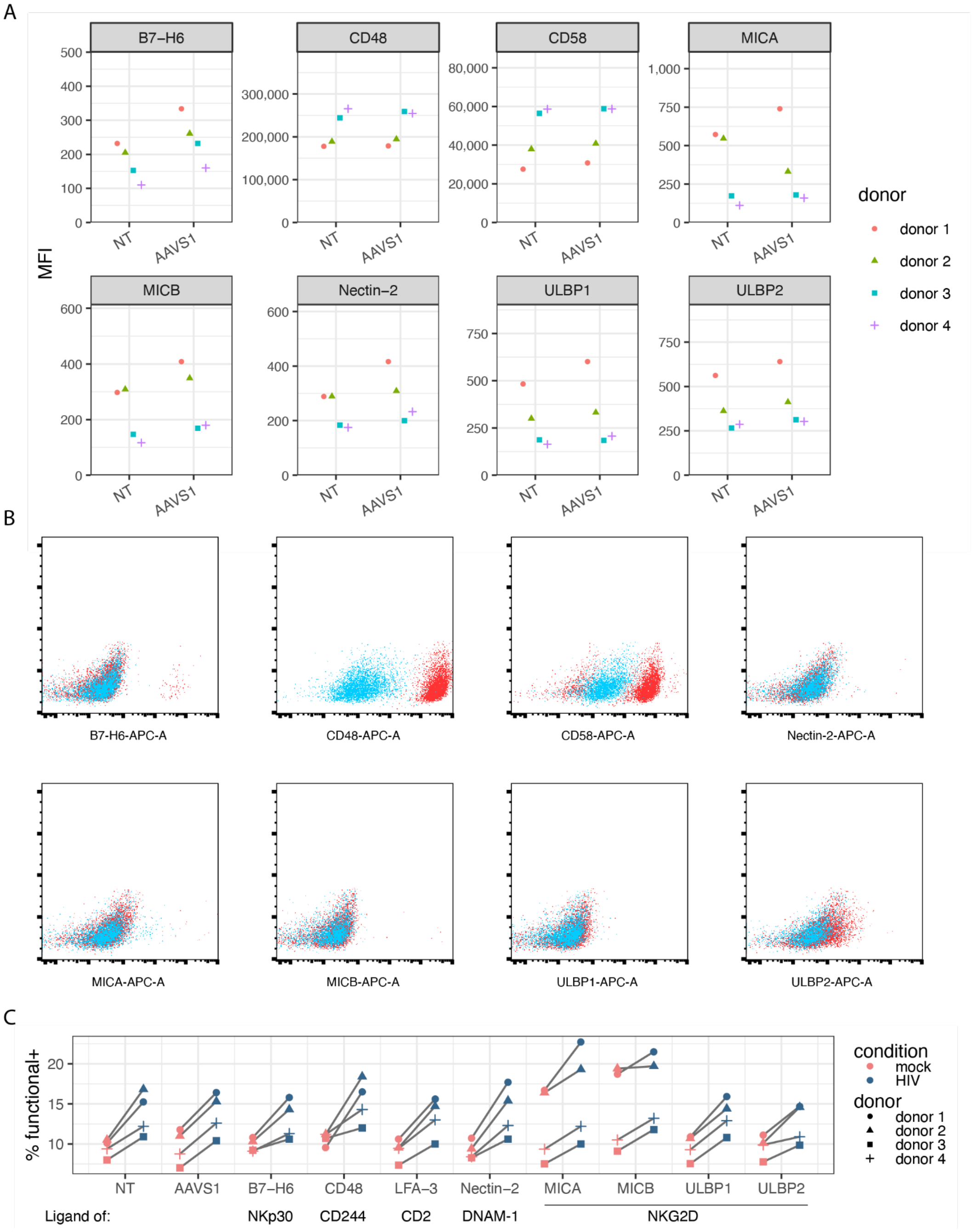


Figure S3: CRISPR knockout of individual genes that encode NK cell ligands in CD4⁺ T cells
**(A)** NK ligand expression in untreated and AAVS1 controls. Mean fluorescence intensity (MFI) of 8 targeted NK cell ligands, in non-treated (NT) and AAVS1 KO (AAVS1) controls. Each donor is indicated by a different symbol. **(B)** Representative flow cytometry plots comparing the expression level of NK cell ligands in $CD4^+$ T cells from donor 2 with CRISPR knockout of each individual ligand (in blue) or without treatment (NT, in red). **(C)** Percentage of $functional^+$ NK cells (cells that were stained positive for any of CD107a, IFN-γ, and TNF-α) indicating NK cell responses to mock- (in pink) and HIV-infected (in blue) $CD4^+$ T cells, in all knockout conditions or NT control after NK cells and $CD4^+$ T cells were co-cultured at effector:target ratio of 1:1. n=4. Each donor is indicated by a different symbol.

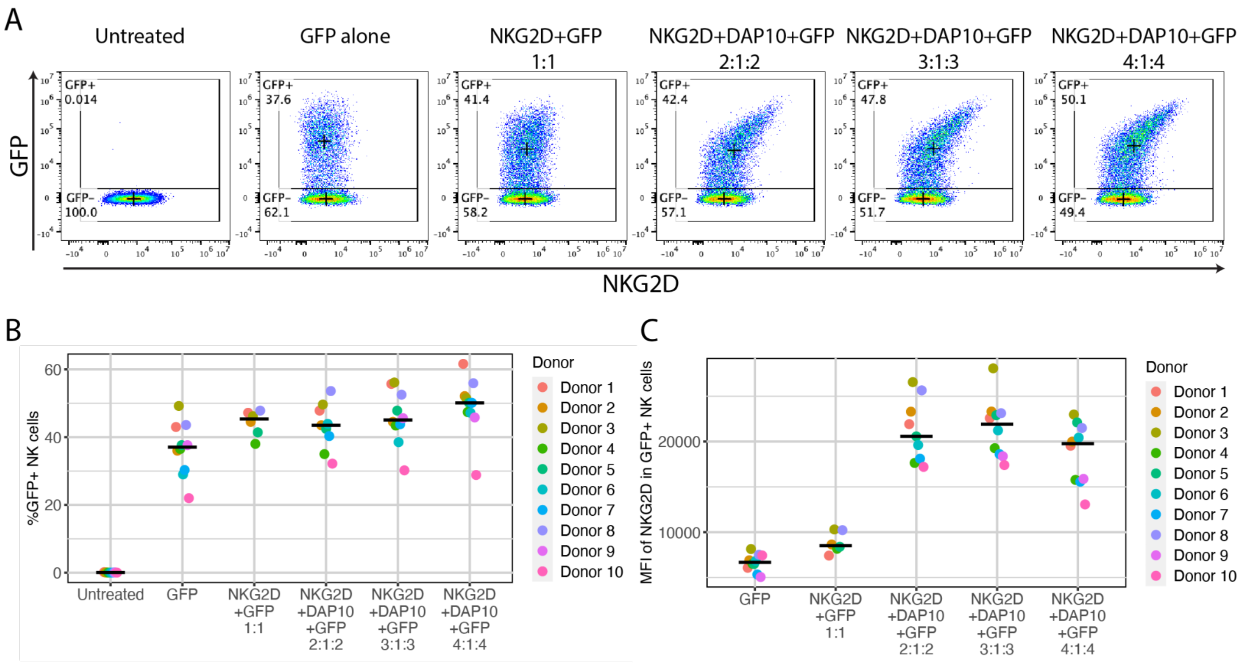


Figure S4. Co-transfection of NKG2D, DAP10, and GFP mRNA in primary NK cells with CART. **(A)** Representative flow cytometry plots indicating the co-expression of GFP and NKG2D in NK cells purified from PBMCs of healthy donors after the NK cells were transfected with the encoding mRNA according to the ratios as indicated on top of each plot. Crosses indicate the median fluorescence intensity of the marker on x and y axes in each gate. **(B)** Percentage of NK cells that express GFP after the cells were transfected with the encoding mRNA according to the ratios labeled on x axis. Bars indicate the median in each group. n=6-10. **(C)** Mean fluorescence intensity (MFI) of anti-human NKG2D-PE signal indicating the expression level of NKG2D in GFP$^{+}$ NK cells after NK cells were transfected with the encoding mRNA according to the ratios labeled on x axis. Bars indicate the median in each group. n=6-10.

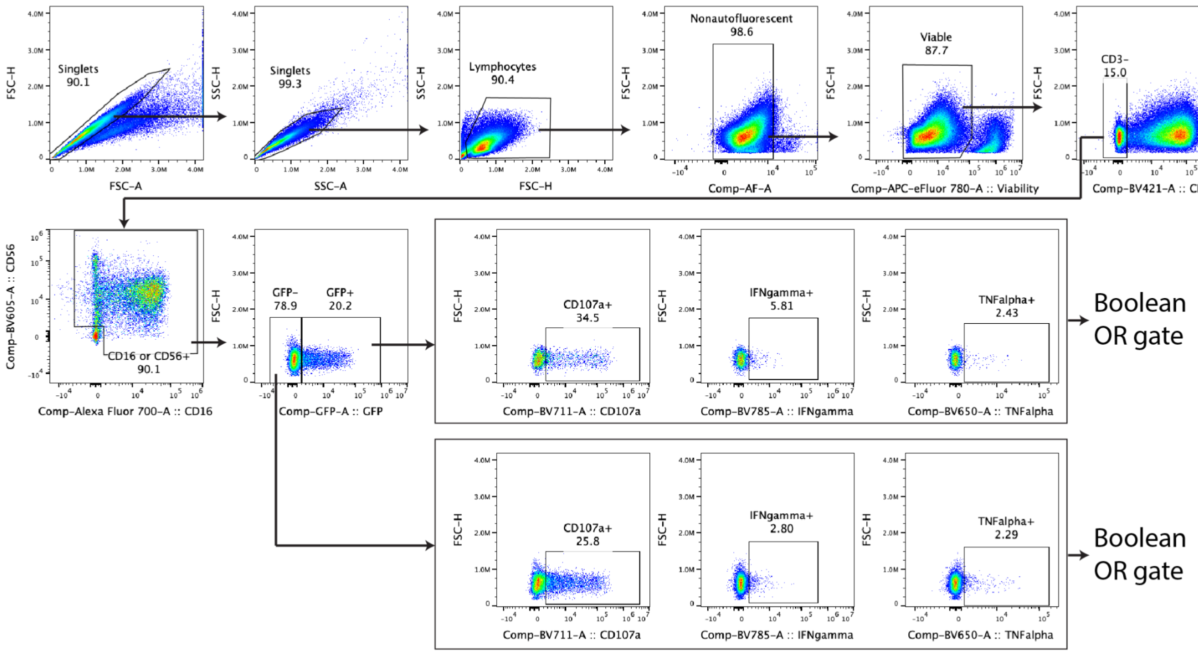


Figure S5. Gating strategy of identifying the transfected ($GFP^+$) and untransfected ($GFP^-$) NK cells in NK/$CD4^+$ T cells co-culture assay as shown in figure 5D.

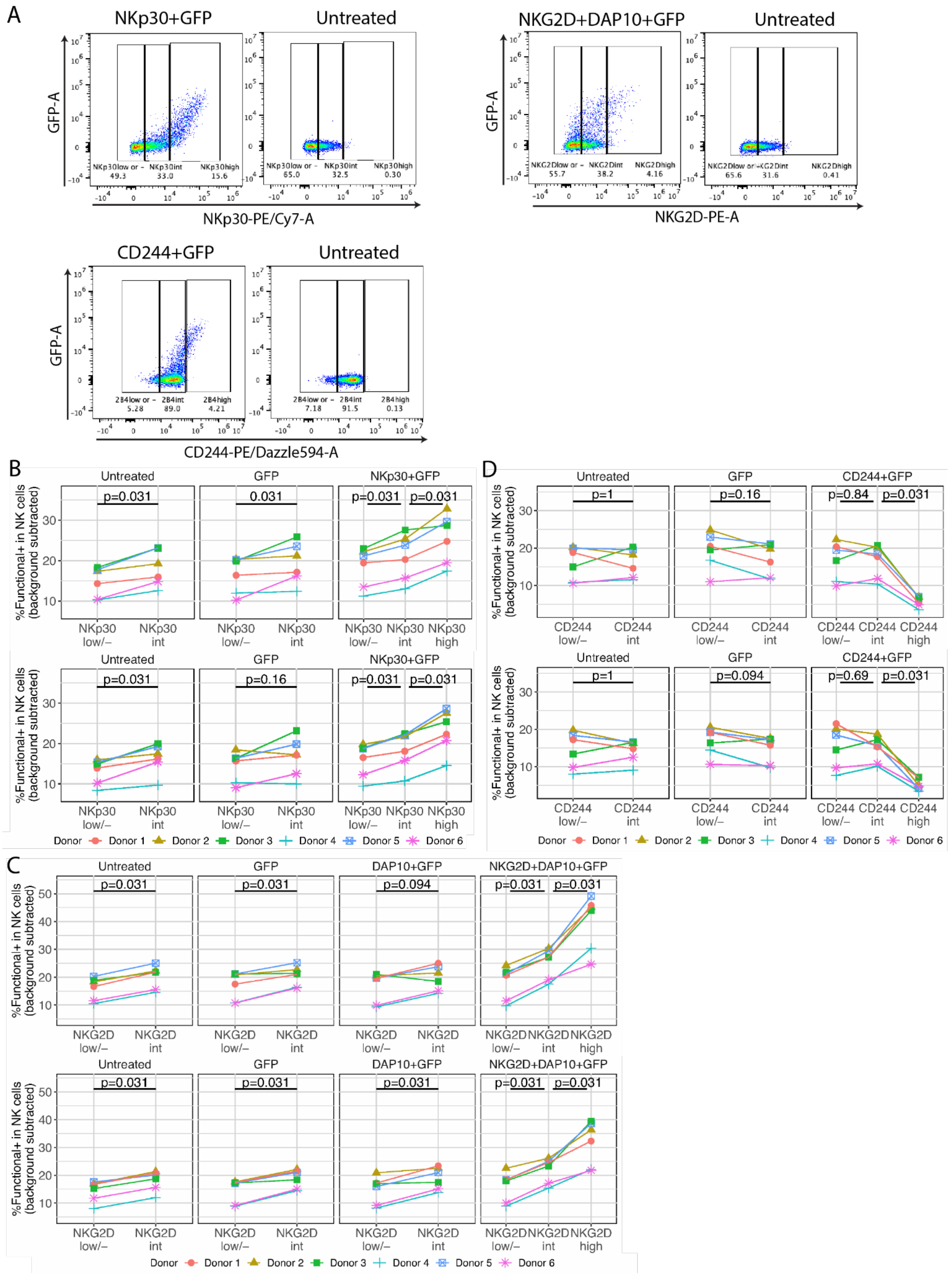
A
NKp30+GFP
Untreated
GFP-A
NKp30low or - 49.3
NKp30int 33.0
NKp30high 15.6
NKp30low or - 65.0
NKp30int 32.5
NKp30high 0.30
NKp30-PE/Cy7-A
NKG2D+DAP10+GFP
Untreated
NKG2Dlow or - 55.7
NKG2Dint 38.2
NKG2Dhigh 4.16
NKG2Dlow or - 65.6
NKG2Dint 31.6
NKG2Dhigh 0.41
NKG2D-PE-A
CD244+GFP
Untreated
2B4low or - 5.28
2B4int 89.0
2B4high 4.21
2B4low or - 7.18
2B4int 91.5
2B4high 0.13
CD244-PE/Dazzle594-A
B
Untreated
GFP
NKp30+GFP
p=0.031
0.031
p=0.031
p=0.031
%Functional+ in NK cells (background subtracted)
NKp30 low/–
NKp30 int
NKp30 high
Untreated
GFP
NKp30+GFP
p=0.031
p=0.16
p=0.031
p=0.031
Donor
Donor 1
Donor 2
Donor 3
Donor 4
Donor 5
Donor 6
D
Untreated
GFP
CD244+GFP
p=1
p=0.16
p=0.84
p=0.031
CD244 low/–
CD244 int
CD244 high
Untreated
GFP
CD244+GFP
p=1
p=0.094
p=0.69
p=0.031
C
Untreated
GFP
DAP10+GFP
NKG2D+DAP10+GFP
p=0.031
p=0.031
p=0.094
p=0.031
p=0.031
NKG2D low/–
NKG2D int
NKG2D high
Untreated
GFP
DAP10+GFP
NKG2D+DAP10+GFP
p=0.031
p=0.031
p=0.031
p=0.031
p=0.031

Figure S6. Correlation between NK cell response to $CD4^+$ T cells and expression level of NKp30, NKG2D and CD244

**(A)** Representative flow cytometry plots indicating the gating strategy to stratify NK cells into populations that express high, intermediate (int), or low/negative level of NKp30, NKG2D, or CD244 in NK cells that were co-transfected with the mRNA as labeled or in untreated NK cells. Percentage of $functional^+$ NK cells (positive for any of CD107a, IFN-ɣ, and TNF-α) in NK cell populations that express high, intermediate (int) or low/negative level of NKp30 **(B)**, NKG2D **(C)**, and CD244 **(D)** after transfected with the mRNA as labeled on top of the panels and co-cultured with autologous HIV-infected $CD4^+$ T cells (top panel of (B),(C) and (D)) and mock-infected $CD4^+$ T cells (bottom panel of (B), (C), and (D)) after subtracting the percentage of $functional^+$ NK cells in NK alone group (background subtraction). n=6 in (B), (C), and (D). Statistical analysis was performed with Wilcoxon signed-rank test (nonparametric paired t-test) in (B), (C) and (D).
